# Microplastics found in high concentrations around Kantamanto, the world’s largest secondhand textile market in Accra, Ghana: A citizen science study

**DOI:** 10.1101/2024.05.13.593950

**Authors:** Dimitri D. Deheyn, Joe Ayesu, The Or Foundation, Branson Skinner

**Author notes:** Contributed equally to the work (Dimitri D. Deheyn, Joe Ayesu). List of personnel of The Or Foundation and citizen scientists who provided valuable help: Patrick Abesiyine Anyebuno, Frank Aboyinga, Emmanuel Akyea, Brown Addoquaye, Bright Anokye, Jacob Ayesu, Bright Ayikpa, Priscilla Danso, Abena Essoun, Emmanuel Ewudzei, Al-Hassan Fattawu, Clémence Faure, Musah Iddrisu, Suleman Issifu, Maxwell Klutse, Freeheart Kordah, Somed Mohammed, Nutifafa Mensah, Katia Osei, Sammy Oteng, Peter Yeboah, and Liz Ricketts and Megan Schuknecht. The raw data from this paper are freely available from the Deheyn Lab or The Or Foundation, in addition to the ones made available already in the Online Resources.

## Abstract

Located in Accra, Ghana, Kantamanto is the world’s largest secondhand resale and upcycling market for clothing and textiles, receiving tons of garments every week exported under HS Code 6309 by countries in the Global North. We looked at the abundance of microplastics and microfibers from textiles and other sources in this dynamic environment. Our primary interest was assessing how microfibers in a textile-rich environment contribute to overall microplastic pollution. Continuous monitoring of air quality using PurpleAir sensors showed large concentrations of airborne microparticles around the market, which were shown to drop with rain events. Water collected from rain contained dozens to hundreds of microfibers and microplastics (roughly in a 1:10 ratio), depending on the intensity of the rain and on whether the rain was preceded by a significant period of dry weather. Microfiber and microplastic counts from airborne samples showed concentrations about >20x to >100x greater, respectively, than those reported from other metropolises around the world. A comparison of concentrations from water in the adjacent Korle Lagoon was similarly striking, with up to 45x more microfibers, and up to 200x more microplastics than what has previously been reported for lagoon waters worldwide. Beyond the microscale, the issue of whole garments as waste was also demonstrated, with a strong similarity found between the global brand garment tags from fashion items discarded in the market and those that were washed up on nearby beaches, highlighting the need for more holistic regulatory frameworks to address textile waste.

## INTRODUCTION

Significant work has been undertaken to establish the environmental footprint of textile manufacturing operations (Madhav et al. 2018; Niinimäki et al. 2020; Grazzini et al. 2021; Farhana et al. 2022; Lim et al. 2022) and to characterize pollution resulting from garment use and maintenance at the primary consumer level (Carr 2017; Carney Almroth et al. 2018; Liu et al. 2019; Nowack et al. 2021; Liu et al. 2023). In contrast, efforts remain largely limited with regards to assessing the environmental impact of garments once they are discarded for secondhand reuse or for end-of-life in a landfill or incinerator. In general, fashion waste originating worldwide ultimately ends up in the Global South, particularly in African countries (Manieson and Ferrero-Regis 2021; UNComtrade 2023; OEC 2024), where the socio-environmental impacts of the waste remain poorly addressed.

Accra (Ghana) is the recipient of tons of fast fashion garments weekly and is home to Kantamanto Market, the largest secondhand textile trade market in the world. The market is adjacent to Korle Lagoon, a major water body found in the city’s center with an outlet that connects to the Gulf of Guinea. The market is connected to the lagoon via a network of small drainage systems and outflow pipes feeding a tributary canal that leads to the lagoon. The area is mostly pedestrian, and the market has over 10,000 stalls, all exposed to some degree of open-air circulation though most are protected under a partial roof. About 15,000,000 garments exported from Global North countries enter the market every week, primarily on Thursdays, in the form of 55 kg bales that the market retailers purchase from importers (The Or Foundation 2022). The retailers work to recommodify the individual items through resale, repair and remanufacturing during the week. As for the undesired or damaged items that go unsold (the majority of which are pre-sorted for reusability prior to arrival in Ghana), they are eventually cast aside, accumulating as waste around the market, with piles of them being burned during the night to avoid excessive build-up, as more space is continuously needed. As such, with Accra’s waste infrastructure unable to handle and safely process the continuously increasing volume of low-quality garments (Niinimäki et al. 2020), the resulting textile waste can easily enter the surrounding ecosystems. This can take place through airborne processes, by burning textiles with microparticles flying into the air, and/or though waterborne processes by dumping garments regarded as waste (full or in pieces) in uncontrolled sites of the lagoon shoreline that are adjacent to the market.

Over the years, the market has grown to become a critical economic center for the region, where an estimated 30,000 people converge every day to work as market traders, upcyclers or to provide auxiliary support. Considering customers, the number of people buzzing through the market goes up to around 60,000 total daily. This large number of people combined with the constant traffic of large volumes of textiles across the market makes Kantamanto a place with potential exposure to microparticles, both microplastics (from packaging material, bags or film wraps) and microfibers (from clothing). Indeed, although the washing and drying of textiles are obvious sources of shed microfibers, another important source is related to wearing them and hanging them outside where they are exposed to various environmental conditions (including high temperatures, humidity, wind and UV) that facilitate their shedding (Carney Almroth et al. 2018; Belzagui et al. 2019; De Falco et al. 2020; Lim et al. 2022; Zhang et al. 2022). Thus, this study offers new evidence on the release of textile microfibers (viz. fiber fragments, with a length greater than the diameter) in a dense urban setting and additional paths through which textile waste can enter the surrounding ecosystems.

The dominant textile materials in Kantamanto are made of cotton but also plastic material, such as polyester, nylon and elastane (and often in blends of two or more of these), particularly given the global proliferation of cheaply made (and quicker to break down) fossil-based fast fashion and the quick turnover between consumption and then disposal as secondhand textiles (Niinimäki et al. 2020). The area is also the recipient of single use plastics like plastic bags, plastic bottles and food containers that litter the market and surroundings.

Knowing all of the above, we hypothesized that the air and water would contain high concentrations of microplastics, and plastic microfibers as well as cotton microfibers. It is also fair to assume that anyone living or working in Kantamanto, or passing through the market, would be exposed to air pollution, and potentially water pollution, in relation to the ambient load of microplastics and microfibers (both synthetic and natural). We also hypothesized that whole garments found as waste on beaches outside the lagoon would originate from the textile dumpsite of the market, which overflows into the lagoon. As such, the garments would show similarities (in terms of brand assemblage and polymer constituent as identifier) to the garments discarded in the market.

Within this context, the research presented here involved a large contingent of citizen scientists who worked to establish whether Kantamanto Market and its main dumpsite nearby generate environmental changes on the surroundings, specifically with regard to the load of microparticles in the air and in lagoon waters. Our effort was cross-disciplinary, performing the best possible environmental sampling with tools and supplies readily available for citizen scientists to monitor the environment in which they live, to address their concerns in terms of environmental and public health, as well as for strategies to develop more sustainable socio-eco-environmental services. This study is the first to establish the close relationship between fast fashion waste and its subsequent release of microplastics and microfibers in the environment, both in the air and water. Our results add to the evidence emphasizing the significant environmental burden disproportionately borne by the communities that have become the de facto stewards of global waste (Clapp 2025).

## METHODS

The study area was the Kantamanto secondhand textile market in Accra, Ghana, the nearby Korle Lagoon and the contiguous beachfront (Fig. 1), which we investigated in May 2023 for microplastic and specifically microfiber concentrations, using waterborne samples as well as airborne samples. Considering the dependence of our analyses on environmental conditions, we used an ATMOS41 ZL6 weather station (https://metergroup.com/products/atmos-41/) placed at the headquarters of The Or Foundation, which is located on the outskirts of the market, to track weather data. The weather station recorded a series of parameters every 15 min., the most important ones used in this study being the time of day when episodes of rain occurred and the amount of rainfall recorded in mm. For microplastic counts, we needed to collect actual volumes of rainwater (expressed in mL), which we did using a Stratus Rain Gauge (https://www.scientificsales.com/6330-Stratus-Rain-Gauge-p/6330.htm) placed next to the ATMOS Station (Online Resource 6). Although the ATMOS Station collected data continuously, the rain gauge collected rainfall water only during time periods that were possible and safe for the citizen scientists to place the gauge in and out of its operating location on the roof. As such, the volumes of rainwater analyzed did not correlate exactly with the precision rainfall measurements of the ATMOS Station due to the difference in the respective range of operating periods. Our study does not intend to be a weather study and for this reason we referred to the amount of rainwater collected in mL, which is the most accurate for our purpose, while disclosing mm of precipitation for reference. Following collection, rainwater was then analyzed for concentration of microplastics and microfibers as described below, based on volumes of mL collected.

**Figure 1.**
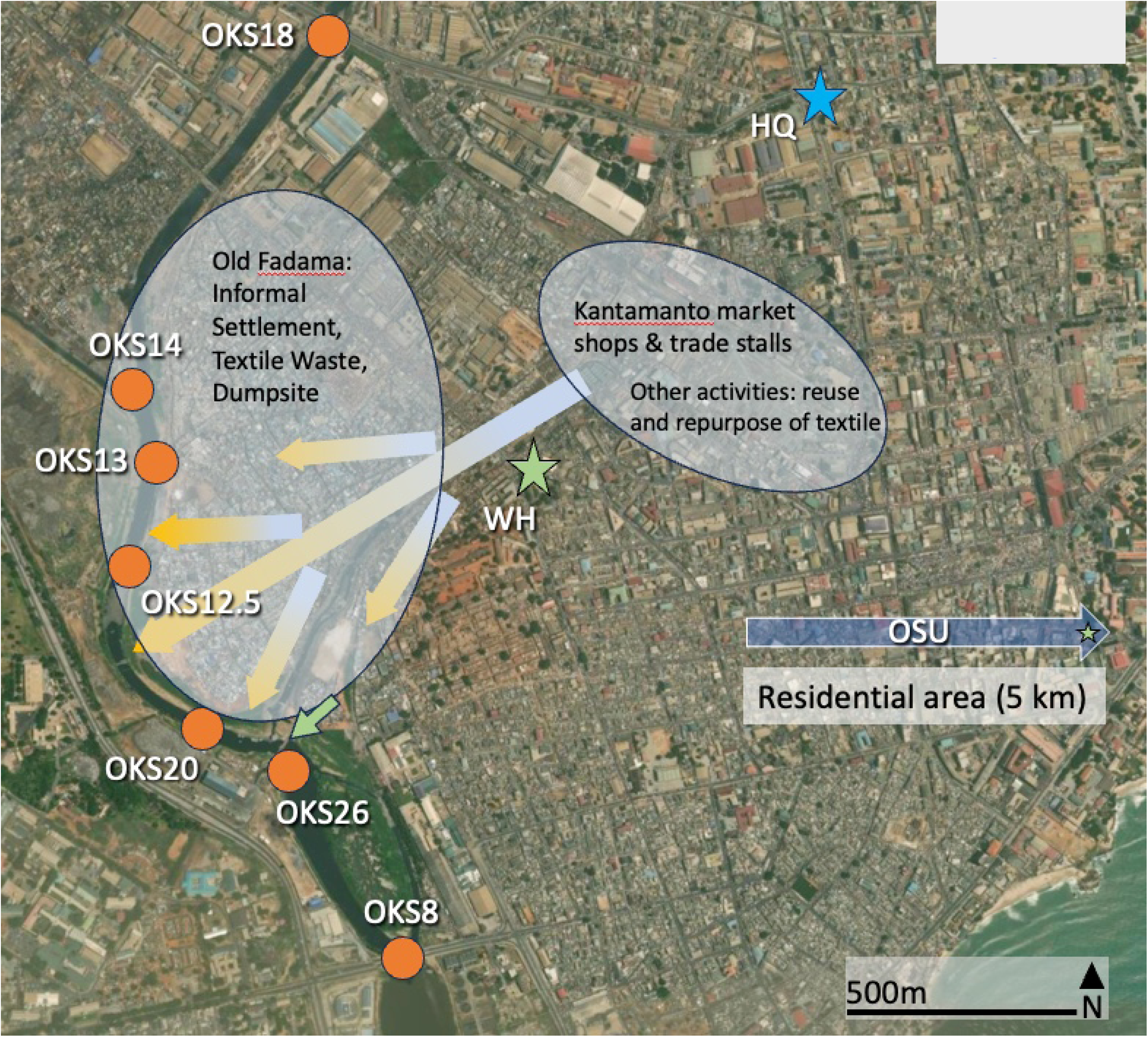
Map (ESRI) of the areas of investigation around Kantamanto Market, indicating the locations for the PurpleAir sensors and the Coriolis air sampler (green and blue stars, respectively), and the water sampling sites in the Korle Lagoon (orange circles). Location of the surface barrier is also visible (green arrow). Main directions of surface water runoff and surface wastewater are also indicated (yellow arrows). HQ: Headquarters of The Or Foundation; WH: Warehouse, large, enclosed space immediately adjacent to the market where textile waste is prepared for reuse; OSU: Residential neighborhood in central Accra, about 5 km from the market.

Sample collection for microplastics and microfibers has become (since 2020) a routine monitoring activity of The Or Foundation (Ghana), which was put together and optimized over the years with the Deheyn lab at Scripps Oceanography (California, USA). The work in the field, and in the lab, is always done with what is readily available locally in a non-academic infrastructure, keeping in mind the possibility of contamination (Chen et al. 2020; Prata et al. 2020; Song et al. 2021; Ma et al. 2024), which is addressed with various precautions and controls. In the field, collectors wear bright yellow safety jackets and bright blue/yellow gloves (the microfibers of which are easy to detect under an imaging microscope in brightfield illumination), and samples are collected standing downstream from the actual collection process. Samples are brought back in closed and cleaned containers to The Or Foundation headquarters. There they are processed in an enclosed laboratory with no windows and no direct exposure to outside air, thus being relatively sealed off from the surrounding activities of the local area. The samples are processed using conventional guidelines against contamination, only using tap water passed through a ceramic clay filter (pore size < 1μm) and limiting direct exposure to air flow (samples are protected by non-sticky parchment paper and aluminum foil when needed). Negative (blank) controls of all the processing steps are performed on a regular basis (on average once a month or when something is changed in the laboratory, e.g. water containers) in order to identify possible airborne and waterborne sources of contamination. This includes running the processing steps of sample preparation as usual but without the sample. Such blanks are also run for each solution used, for each step (e.g., sitting in an oven, fridge or under the microscope), considering the same volumes and times as for regular sample processing. The concentrations of microparticles found in this range of controls are considered “negative” because they are not associated with the sample and are removed from the data presented. For this paper we focused on the month of May 2023, because this was a month of heavy rain during which all of our sensors worked without interruption (viz. there was no power outage from the city grid).

*On-line monitoring of airborne microparticles.* A key air quality monitoring sensor is the PurpleAir, a small “accessible-to-everyone” instrument that detects concentrations of particulate matter in air at a given time point (Ko et al. 2020), which has been used widely around the world by people (technically citizen scientists) interested in knowing the quality of the air they breathe (note that the PurpleAir does not collect or trap the airborne microparticles). The monitoring is continuous and the data (often made publicly available in real-time) characterizes the air quality in a few categories (www.PurpleAir.com), which are in line with the US EPA Clean Air Act quality standards (Barkjohn et al. 2021; Jaffe et al. 2023).

In this study, PurpleAir PA-II-Flex air quality sensors were used to monitor concentrations of PM2.5 (smaller than 2.5 microns) and PM10.0 (smaller than 10 microns) particles expressed as mass concentration (μg/m^3^) in the air, which is the conventional unit to report air quality under the standards of the US EPA Clean Air Act. This mass concentration output, however, originates from raw numbers of particles that are transformed into mass considering specific algorithms (see www.PurpleAir.com) and assumptions about the particles (mostly that they are pollen, dust, and/or black carbon from improper fuel combustion). This mass concentration could therefore be biased under conditions with heavy concentrations of airborne microplastics (which we assumed would be the case around Kantamanto Market, considering its substantial volume of synthetic clothing). As such, we also used the raw data of these sensors, using the PM2.5 and PM10.0 particle counts directly, which are expressed in dL by the sensor (and expressed in L here). These particle count concentrations can therefore be more reliable to compare to the data obtained from the Coriolis® air sampler (see section below).

One PurpleAir sensor was located next to the ATMOS41 weather station, in the area near the market corresponding to the upper deck (4^th^ floor, about 12 m above street level) of The Or Foundation headquarters (Fig. 1). Another sensor was positioned on a metal pole at either 5 m or 10 m height and moved around for measurements in different locations, including in the heart of the market. Measurements were also taken 5 km away from the market, in Osu, a residential area of Accra. Data from the PurpleAir were processed according to the protocol recommended by the vendor. (The raw data, which was collected every 2 min, was organized in a spreadsheet and curated for any missing data, such as for when the instrument was saturated e.g., or for when power supply instability would make it miss some measurements). Although PurpleAir sensors measure the concentration of particles within their range size, they do not identify what the particles are in terms of their chemical makeup. As such, they do not identify whether particles have a natural basis (dust, pollen vs black carbon) or are purely synthetic, like microplastics (e.g., such as tire shred particles or microfibers from textile fabrics). For this reason, airborne particles had to be collected for infrared fingerprint analysis to identify their chemical nature.

*Airborne sample collection.* A Coriolis Compact portable air sampler instrument (Bertin Technologies) was used to collect airborne microparticles between 0.5 and 10 μm from the same locations as the PurpleAir sensors. (Note that the manufacturer has validated the optimal efficiency of the Coriolis for this size range of microparticles, which does not exclude that larger particles can also be collected, since there is no physical size exclusion mechanism in place but rather pelleting of particles from generated cyclonic air flow conditions in the instrument; https://www.bertin-technologies.com/products-range/air-samplers/). Because the instrument collects the microparticles, it cannot be run 24/7 like the PurpleAir, which is used for monitoring the microparticles in the air without collecting them and knowing what they are. Thus, the PurpleAir and the Coriolis instruments appear to complete complementary functions, which we took advantage of in this study. Here, the Coriolis was run for 2 hrs each time at each location, with a vacuum rate of 50 L/min as indicated by the manufacturer specifications. The particles that were collected in the 10 mL cone receiver of the instrument were retrieved according to the following process (given that MilliQ water is not readily available): Tap water filtered through ceramic clay filters with pore size <1 µm was added to the cone, shaken using a Vortex, and poured into a 50 mL Falcon tube. The process was repeated five times to make sure all the particles were transferred.

*Waterborne sample collection from the lagoon.* Water was collected from 7 sites (named OKS8 through OKS18) along the lagoon, in order to have samples upstream, running next to, and downstream from the dumpsite that receives significant quantities of textile waste leaving Kantamanto Market (Fig. 1). A permanent cement beam structure (aka surface barrier) crosses the width of the lagoon between OKS20 and OKS26. This surface barrier is about 40 m long and about 1.5 m tall, and the watermark reaches about half-way up the height of the barrier. As such, the surface barrier stops floating waste from moving further down the lagoon and causes items to build up in massive amounts upstream, to the point where the water surface can barely be seen in OKS20. In contrast, OKS26 (located just downstream from the surface barrier) has much less floating waste since it receives the water that flows under the surface barrier, from about 0.75 m below the water surface. The surface barrier thus does not prevent heavier waste (sub-surface, midwater or tumbling on the seafloor) from flowing under the structure, downstream in the lagoon and into the ocean.

Considering the poor quality of the water in the lagoon, which is due in part to raw sewage input (Nixon et al. 2007; Koppelaar et al. 2018; Owusu et al. 2023), surface water collection was performed using an extended pole. The pole was equipped with a dedicated 5 L bucket made of a clean, bright blue plastic easily identifiable under microscope imaging (for blank background). This way, the water collection was safe for the operator. Water was collected with the bucket, the bucket was rinsed and emptied on the ground, and water was collected again 2 separate times at the same site. Each time two 50 mL Falcon tubes (N=4 total) were rinsed three times with lagoon water from the bucket, shaken vigorously. emptied, and rinsed/emptied again two times, and then filled with water from the bucket for the fourth time (for analysis), ensuring that the tubes were submerged sub-surface during collection (to limit any possibility of airborne contamination). The tubes were capped, labelled and transported in a cooler back to the laboratory.

The water collection method put in place in this study, based on collecting 50 mL Falcon tubes of liquid, is unique in allowing samples to be collected from almost anywhere, and by almost anyone. This is powerful for gathering a large number of samples (and replicates) from places that are difficult to access in time or space. This methodology therefore leads to numbers of microplastics and microfibers for every 50mL of sample collected, which is the concentration that will be expressed throughout the study (counts/50mL), since expressing the data per L (as is common) could induce a biased extrapolation. Our data can be compared to microplastic concentrations found in the literature, yet keeping in mind the difference in methodology and the difference in sample size and processing. Indeed, most studies focusing on microplastics use the Manta Net to trawl large volumes of water, thus results are expressed in unitary m^3^ or L, and only available from coastal areas where boats can operate. The Manta Net usually has a mesh size of 100-500 μm, thus collecting larger microplastics, rather than the smaller microplastics (<100 μm), as we do in this study. This difference in methodology relative to coastal baseline studies on microplastics and the difference in size range of microplastics collected, needs to be kept at the forefront when reading this paper, with the difference in units a reminder of these fundamental differences.

*Samples processing for microplastic and microfiber counts*. Samples were brought back to the laboratory where they were processed for microplastic and microfiber counts following a routine protocol used in the Deheyn Lab at Scripps Oceanography, (UC San Diego, USA) and implemented at The Or Foundation (see Online Resources). This involved filtering the samples on glass fiber filters (0.6 μm mesh size to retain very small microplastics) and imaging the dried filters under a stereomicroscope. The imaging was done in bright field as well as epifluorescence (ex. 360-380 nm, em. >415 nm Long Pass filter; see Online Resource 1-2 for details). Microplastics and microfibers were counted by at least 3 individuals (and up to 6 individuals) using both the bright field image and the epifluorescence images, since they showed different items (Online Resources 1-2).

The epifluorescence property of materials of different colors was addressed in the Deheyn Lab, using reference fabric material provided by industry (Lenzing AG and Archroma). We showed that synthetic white polymers (e.g., polyester, elastane, nylon) displayed vivid blue/cyan blue fluorescence, while their colored counterparts showed no visible fluorescence under the same settings. When mixed with plant debris or phytoplankton, the blue/cyan blue fluorescence was visible together with the red fluorescence of chlorophyll, which we expected was the most representative of what we would find on our filters. As for the nature-based polymers (e.g., cotton, lyocell), they had no visible fluorescence, whether white or colored, under the settings used for viewing their synthetic equivalent (Online Resource 1).

The intent was not to perform an exhaustive study on the effect of color dyes on the fluorescence but to demonstrate using key polymers and representative colors that our combination of bright field and epifluorescence counting was comprehensive enough to sort microparticles into four categories: (1) Colored microparticles from a variety of sources, including textiles, degrading plastic items, or even just silt grains (bright field counts); (2) Microparticles from natural debris rich in chlorophyll, whether decaying plants, filamentous algae or cyanobacteria (red fluorescence counts); (3) Synthetic microplastics (blue fluorescence dots, flakes or particles); and (4) Synthetic microfibers (blue fluorescence fibers) (Online Resource 2). This combination of bright field and epifluorescence imaging thus allowed us to broadly cover different situations of the life of plastic. For example, an unweathered piece of colored plastic will be seen in bright field but not necessarily in fluorescence, since dyes could affect the innate synthetic polymer fluorescence by shifting its intensity and/or color (Grieve et al. 2005; Biermann 2007). In contrast, weathered pieces of plastic, invisible in bright field, will emit bright blue fluorescence. As for red fluorescent items, they will likely represent a rich content in chlorophyll, which could be seen in bright field as well since they display a green color, unless the red fluorescence originates from a synthetic dye, which would likely be seen in red in bright field (Online Resource 2).

The synthetic microfiber counts, which represented our main interest, are discussed in the Results section. (The counts from the other analyses, which are from bright field, microparticles red fluorescence, and synthetic blue, fluorescent microparticles are available as Online Resources. They show the same trends as the synthetic microfibers but in different ranges. Nonetheless, similar conclusions can be drawn from this data.)

A subset of samples (35 items) was processed for polymer chemical identification using Quantum Cascade Laser InfraRed (QCL-IR) spectral imaging, using the Spero® IR system (DLS DayLightSolutions Inc.) (Primpke et al. 2020), which is available at the Deheyn Lab (see Online Resource for detailed methodology). The infrared hyperspectral cubes (950 to 1800 cm^-1^) generated from each sample were compared to those of 23 reference standards for plastics (PolymerKit 1.0; Hawaii Pacific University) and 7 raw textile fibers that were provided by industry collaborators at Lenzing AG and Archroma. The infrared fingerprint spectral data for three common polymers standards were overlapped over the spectral data collected from some microparticles to illustrate our polymer assignment (see Online Resource 3).

*Textile waste by brand tag*. We assessed the relationship between the textile waste as whole garments collected directly from retailers in Kantamanto Market and the macro textile waste pollution found in the lagoon and at the beaches flanking the opening of the lagoon. For this, we compared the brand tags from garments found at the market (172 garments) with those found at the beach (128 garments) (see Online Resource for details). We expected the assembly of brands from the beach textiles to reflect the ones from the market, since we presumed that the discarded garments somehow (through the lagoon flow) made their way to the ocean, where the tides then pushed them back on the local beaches. We then correlated the two sets of tags using a Pearson correlation coefficient to assess their potential relationship, viz. the market, via the dumpsite, being the source of the textile waste found on the beach. In addition, we recorded the polymer textile fiber composition of the garments (relative content of cotton and polyester in each garment, for example) as indicated in the product tag. In that case, we expected their polymer composition to reflect the one of microfibers found in the airways and waterways.

*Statistical analysis.* This study aims to provide a snapshot of the concentration of microplastics and microfibers in the dynamic environment of the world’s largest second-hand textile market. A diversity of variables is considered, from textile-related merchandising activities to textile burning, in addition to weather variables and to the schedule of the crowd coming in and out of the market. Our analyses therefore mainly rely on descriptive statistics using box plot displays, with data curing performed in Excel (Microsoft Corp) and figures made in Delta Graph (Red Rock Software). Although our intentions were not to test the particular level of significance of variables, we performed the research with as many replicates and independent controls as possible, in order to provide a robust assessment of effects and trends. The only “experimental” assessment we did, which deserved statistical assessment, was when we compared the assembly of brand tags from the market textile waste (which we manually pulled out of the market) versus the one from the beach textile waste (which we dug out from the sand). We therefore ranked the brands from the most abundant to the least abundant based on the number of tags found for each brand and used Excel to run a Spearman ρ correlation coefficient analysis considering a significance level of α<0.05.

## RESULTS

The results showed that the clothing from global sources sold into Kantamanto Market and discarded from the market as waste is a source of microparticles, but that the release and distribution of these microparticles can show large discrepancies, based mostly on market activities and weather (rain), which are variables both in time and space. The synthetic microfiber counts we report here always showed high values compared to their blank controls equivalent for background contamination. In general, our various controls accounted for <2 % of counts compared to those from processed samples (Online Resource 4-8).

Of the microparticles collected via air sampling (Coriolis analysis) and counted by microscope imaging, we show here the data for the synthetic microfibers and microplastics identified in epifluorescence, while the other imaging data is shown in Online Resource 5-8. Overall, all the imaging data followed the same trend but in different ranges of counts. In general, bright field counts were always the lowest and below one hundred, followed by red fluorescence counts and the blue fluorescence microfiber counts (both about 10x greater than the bright field counts) and blue fluorescence microparticle counts (about 10x greater than the blue fluorescence microfiber counts). Broadly, this indicates our samples contained a sustained level of recently released microparticles: a large amount of natural microparticles and synthetic microfibers and a much greater amount of microplastics (Online Resource 5-8).

*PurpleAir data were up to 5x greater within the market*. PurpleAir sensors allowed us to collect data over a continuous 24 hr period and to measure both mass concentration and raw counts of particles (N=720 data points each). The PurpleAir sensors across the different areas of town and at the different height (5 m, 10 m) all showed similar mass concentration data in terms of the relative abundance of PM10 and PM2.5 particles (as expected since PM10 data encompass PM2.5 data), which systematically showed PM10 mass concentration data up to 125% greater than that of PM2.5 (Online Resource 4A). The most striking difference across sites was regarding the variability of the data, which was more variable in the market than off market, and which thus translated into a greater difference between the means and the medians for each of these sites (Online Resource 4A). Over the course of May 29, 2023, which was a day with no rain (last rain event was about 40 hrs prior) PM2.5 in the air at a given timepoint ranged from 3 to 356 μg/m^3^, while those of PM10 ranged from 4 to 372 μg/m^3^. The mass concentration was systematically greater for the sensors placed in the market, especially for the one at 10 m height (PM10 up to 372 μg/m^3^), which was slightly greater than the one at 5 m height (PM10 up to 332 μg/m^3^). The mass concentration decreased when moving away from the market, whether at its outskirts at The Or Foundation headquarters at 12 m (up to 103 μg/m^3^) or in the residential area control at 2 m (with up to 94 μg/m^3^) (Online Resource 4A). The corresponding data with respect to microparticle count concentration (raw data generated by the PurpleAir) showed similar characteristics, with greater count concentration and greater variability for sites within the market (Online Resource 4B). This was especially true for data of PM2.5 particles, which was up to 3,467 counts/L in the market and up to 912 counts/L off the market. The particle count concentration for PM10 were lower than for PM2.5, as expected, reaching 172 counts/L in the market and up to 226 counts/L off the market, indicating that larger microparticles can also be found off market (Online Resource 4B), but whether they originate from the market remains to be determined (see assessment from the section on the Coriolis data below).

The trend in the data, showing greater values in the market relative to the outskirts and residential areas, and, within the market, higher concentrations at 10 m relative to 5 m, was observed throughout our study, yet was highly variable from day to day and within a given day. This was dependent on the activities at the market (most active from 05:00 until 18:00 every day but starting as early as 02:00 for the vendors) with possible burning of waste in locations immediately surrounding the market occurring overnight. Vehicle traffic on the market periphery and foot traffic across its web of stalls could also affect the PurpleAir data, but also, and most importantly perhaps, the weather, especially in relation to wind and rain. Indeed, data collected during days when rain occurred decreased sharply as soon as rain started before increasing again gradually after the rain stopped (Fig. 2). Microplastic particles can make up a portion of the overall PM10 and PM2.5 data, and therefore are a component of the air we breathe, with potential impacts on human health. It was therefore critical for us to look at the actual content of microplastics and microfibers for both airborne and rainwater samples.

**Figure 2.**
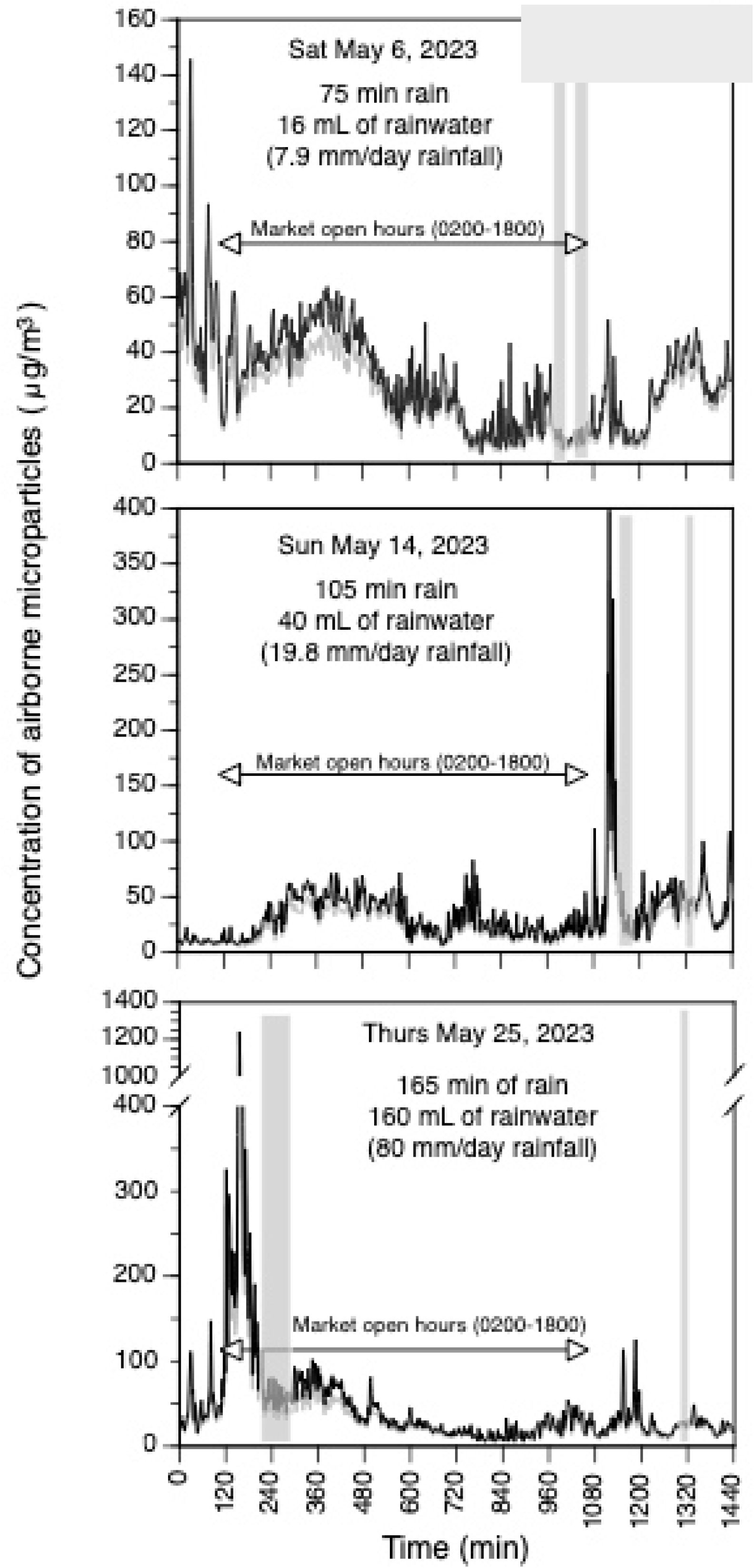
PM2.5 (grey line) and PM10 (black line) particles mass concentration (μg/m^3^) from the PurpleAir data recorded every 2 min during a 24 hr period starting at midnight, for 3 different days with distinct intensity of rain events (date, duration of rain and volume of rainwater collected are indicated). Period of rain (gray shade) and volume of rainfall was provided by ATMOS41 weather station. Period of market activity is indicated on each panel. Of note: Thursdays are when new bales of garments are delivered to the market to be opened; the garments inside are sorted on Wednesdays and Saturdays.

*Airborne microparticles were dominated by synthetic textile microfibers*. Data from the Coriolis air sampler showed relatively high concentrations of microfibers across all sites of investigation within the market, ranging from about 20 at HQ to about 180 microfibers per 6,000 L in the market at 10 m height, equivalent to 0.004 to 0.029 microfibers/L (Fig. 3). In general, we found that the microfiber concentrations were greater for the samples analyzed at 10 m height compared to the 5 m analysis (even within a ventilated warehouse); this was particularly evident for the samples taken in the open-air area of the market (Fig. 3). Measurements in the market were about 2x to 5x greater than the ones made 1 km away, at the outskirts of the market (Fig. 3), which correlated with the data found with the PurpleAir. Greater concentrations of microfibers were also observed in bright field conditions, indicating that these microfibers were not weathered and thus likely originating from the area where the air sample itself was collected (see BF MFs; Online Resource 5). We also found greater concentrations of microplastic particles at 10 m in the market compared to the same location at 5 m. We assumed that such greater concentrations of microplastic particles at 10 m were mostly originating from textiles as well considering that their values were also high in the warehouse where textiles are shredded for upcycling activity, while lower for HQ where textile work is not performed. (see UV MPs, Online Resource 5). These microplastic particles were in much greater concentrations than their synthetic microfiber equivalent, which correlated well with the trend observed for the PurpleAir data where PM2.5 count concentrations were greater than the ones for PM10.

**Figure 3.**
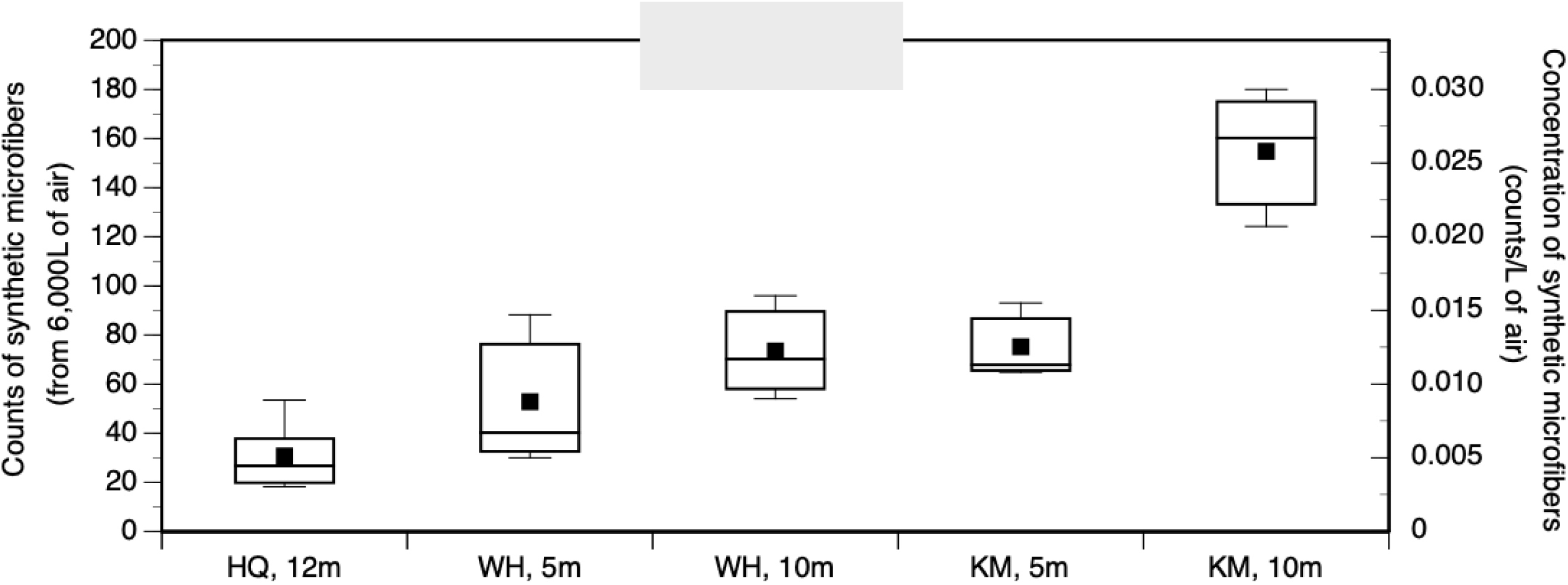
Airborne synthetic microfiber concentrations (box plots; N=9) across the various sites of investigation showing that the values were greater in the market and, within the market, greater at 10 m compared to 5 m. Data were collected with the Coriolis air sampler for 2 hrs each time, thus representing concentrations of “counts/6,000 L of air”. HQ: Headquarters of The Or Foundation; WH: Warehouse; KM: Kantamanto Market.

*Rainwater and lagoon samples contained large concentrations of synthetic textile microfibers.* Rainfall was highly variable and characteristic of a tropical weather regime, thus more often with heavy “showers” scattered in space and time across Accra. We recorded rainfall from 1.5 mm/day to 80 mm/day. Rainwater samples that were collected from the rainfall ranged from a few mL to a couple hundred mL. Analyses showed that the rainwater contained synthetic microfibers (Table 1; Online Resource 6-7) within a range of concentrations (counts/50 mL) that varied with the duration of the rain event, with greater concentrations of synthetic microfibers (up to 300/50 mL) found in rainfall from shorter rain events. Indeed, in situations where we collected over 40 mL of rainwater, which equate to about 19.8 mm of rainfall under weather standards, the concentration of synthetic microfibers decreased sharply (Fig. 4A). In addition, the concentration of synthetic microfibers found in rainwater changed based on when the last rain event occurred, and as such, the greater the extent of dry weather between two rain events, the greater the concentration of synthetic microfibers found in the rainwater. This was especially observed after 3 days of dry weather (Fig. 4B).

**Figure 4.**
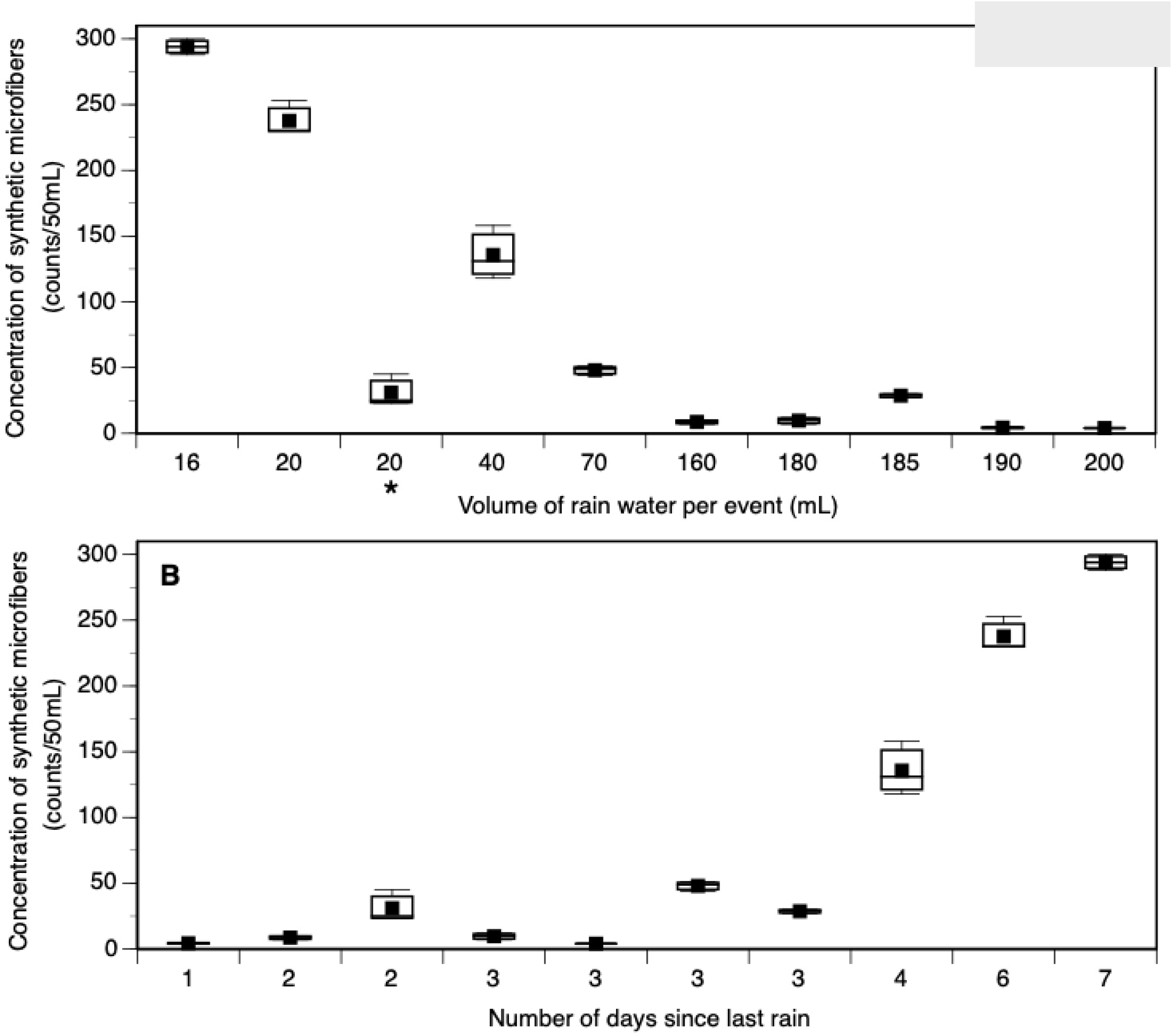
Rainwater microparticle count concentrations (box plots; N=9) collected with the Stratus Rain Gauge (with rain event volumes validated by an ATMOS41 weather station) set on the roof of The Or Foundation headquarters. **A.** Concentrations of synthetic microfibers from rainwater samples that correspond to different rain events, thus each with their own volumes (on the X axis). Data counts on the Y axis are expressed per set volume of 50 mL. The concentration of synthetic microfibers was high for short rain events (up to 2 hrs long) and decreased steadily as rainfall got heavier and/or lasted longer. *These samples were collected the same day as the 185 mL rain event, but just a few hours earlier. This shows that the concentration of synthetic microfibers in volume of rainwater depends on precedent episodes of rain. **B.** Concentrations of synthetic microfibers for rain events expressed in relation to the time (number of days) since the last rain event. Data show that the concentration of synthetic microfibers increases greatly in the rainwater after 4 days with no rain. Data counts are expressed per set volume of 50 mL.

**Table 1.** Minimum (Min), Maximum (Max), Mean and Median values for the counts of synthetic microfibers and microplastics (both found with native blue fluorescence) from samples of air (in 6,000 L), rainwater (in variable volumes), and Korle Lagoon water (in 50 mL set volume sampling standard of our methodology). For all counts, N=9, except for the airborne sampling counts (N=3). Airborne microparticle samples originated from 6,000 L of processed air, and were thus expressed in concentration of “counts/6,000 L”, whereas lagoon water samples were expressed in “counts/50 mL”. Data on concentrations for unweathered microparticles (bright field microscope imaging) and natural microparticles (red fluorescence) can be found in Online Resource 9. <u>Abbreviations.</u> HQ: Headquarters of The Or Foundation (roof) at two different times (a,b); WH: Warehouse space adjacent to the market at two different heights (5 m, 10 m); KM Kantamanto Market (center) at two different heights (5 m, 10 m).

|  |  | Synthetic Microfibers |  |  |  | Microplastics |  |  |  |
| --- | --- | --- | --- | --- | --- | --- | --- | --- | --- |
|  |  | Min | Max | Mean | Median | Min | Max | Mean | Median |
| Airborne particles<br>(counts/6,000 L) | HQa | 8 | 22 | 14 | 11 | 311 | 485 | 414 | 446 |
|  | HQb | 7 | 15 | 11 | 10 | 252 | 290 | 275 | 282 |
|  | WH – 5m | 12 | 35 | 21 | 16 | 1034 | 1700 | 1454 | 1629 |
|  | WH – 10m | 27 | 48 | 37 | 35 | 1348 | 1855 | 1636 | 1704 |
|  | KM – 5m | 52 | 74 | 60 | 54 | 1170 | 1400 | 1290 | 1300 |
|  | KM – 10m | 99 | 144 | 124 | 128 | 1195 | 2930 | 1790 | 1244 |
| Rainwater (HQ) | 16 mL | 92 | 96 | 94 | 94 | 946 | 1020 | 971 | 947 |
|  | 20 mL | 92 | 101 | 95 | 92 | 1358 | 1900 | 1600 | 1541 |
|  | 160 mL | 23 | 32 | 28 | 30 | 385 | 663 | 486 | 411 |
|  | 190 mL | 14 | 18 | 16 | 15 | 421 | 661 | 515 | 462 |
|  | 180 mL | 25 | 43 | 35 | 36 | 3149 | 3601 | 3331 | 3243 |
|  | 200 mL | 14 | 16 | 15 | 14 | 1365 | 1484 | 1427 | 1433 |
|  | 185 mL | 100 | 112 | 107 | 108 | 972 | 1264 | 1082 | 1009 |
|  | 70 mL | 62 | 71 | 67 | 69 | 738 | 1092 | 949 | 1017 |
|  | 40 mL | 94 | 126 | 108 | 105 | 1181 | 1479 | 1370 | 1449 |
|  | 20 mL | 9 | 18 | 12 | 10 | 273 | 285 | 279 | 279 |
| Korle lagoon sites<br>(counts/50 mL) | OKS18 | 3 | 11 | 7 | 6 | 125 | 253 | 187 | 174 |
|  | OKS14 | 4 | 10 | 7 | 7 | 251 | 489 | 328 | 269 |
|  | OKS13 | 5 | 46 | 24 | 22 | 293 | 600 | 443 | 435 |
|  | OKS12.5 | 9 | 18 | 13 | 12 | 208 | 532 | 350 | 395 |
|  | OKS20 | 53 | 81 | 68 | 68 | 246 | 485 | 357 | 364 |
|  | OKS26 | 7 | 15 | 11 | 12 | 74 | 284 | 194 | 182 |
|  | OKS8 | 6 | 19 | 13 | 13 | 42 | 267 | 170 | 165 |

Water samples from Korle Lagoon contained large concentrations of synthetic microfibers, ranging from 2 to 80 counts/50 mL (Table 1; Online Resource 8-9), which were greater when closer to the dumpsite and market compared to the locations that were further away (Fig. 5). In addition, and as expected, the concentrations of synthetic microfibers peaked at OKS20 (between 60 and 80 counts/50 mL, where surface waste is stopped from flowing downstream by the surface barrier crossing the lagoon (Fig. 5). The data was not as clear for the concentration of microfibers identified in bright field, suggesting that the microfibers in the water were mostly weathered and not originating directly from airborne microfibers recently generated in the market (see BF MFs, Online Resource 8). As for microplastic particles, they too increased in concentration in the lagoon areas facing the market (up to 550 particles/50 mL), up to where the surface barrier crosses the lagoon.

**Figure 5.**
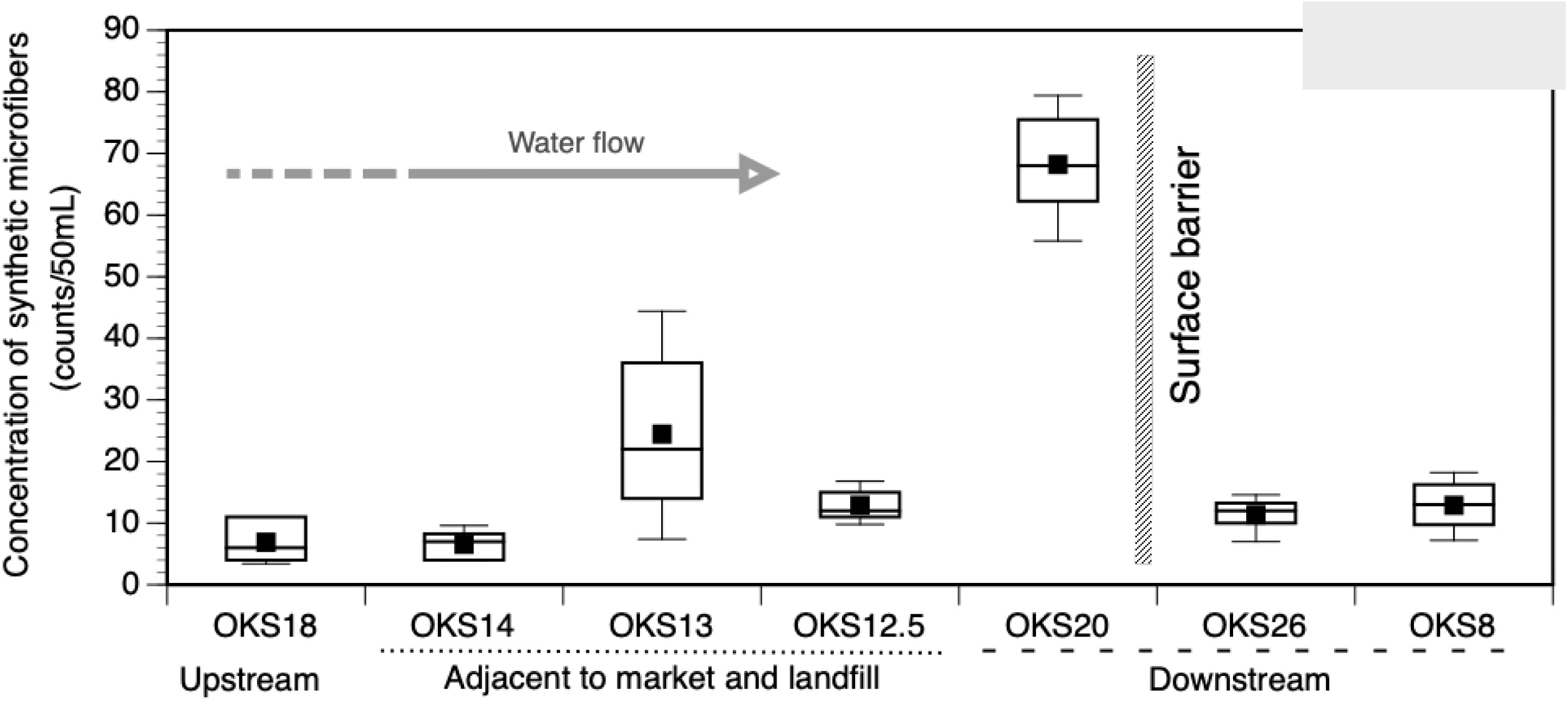
Concentrations (counts/50 mL) of synthetic microfiber (box plots; N=9) from Korle Lagoon water showing that the site directly flanking the market in terms of runoff (OKS13) has the greatest concentrations compare to upstream/downstream of the market, this until the flow of the lagoon water is slowed down by a surface barrier (OKS20), where floating debris and microparticles accumulated. All data counts are expressed per set volume of 50 mL of water collected.

*Chemical composition of waterborne microparticles reflects that of garments from the market*. Infrared fingerprint analyses (see Online Resource 3 for matching of some spectra with standards) showed that waterborne microparticles represented a variety of polymers, the most dominant ones being cotton (29%), polypropylene (26%), polyester (14%), nylon (11%), and polyethylene (9%). Garments in the market reflect this pattern of polymer ratio as well (Fig. 6), with 40% of the garments made primarily of cotton, 13% of polyester and 10% of nylon. As for polypropylene and polyethylene, these polymers are mostly used in plastic bags (used as single-use water containers and takeaway bags) and single-use water bottles, respectively, which are also found spread all over the market and lagoon where they are being weathered and breaking down.

**Figure 6.**
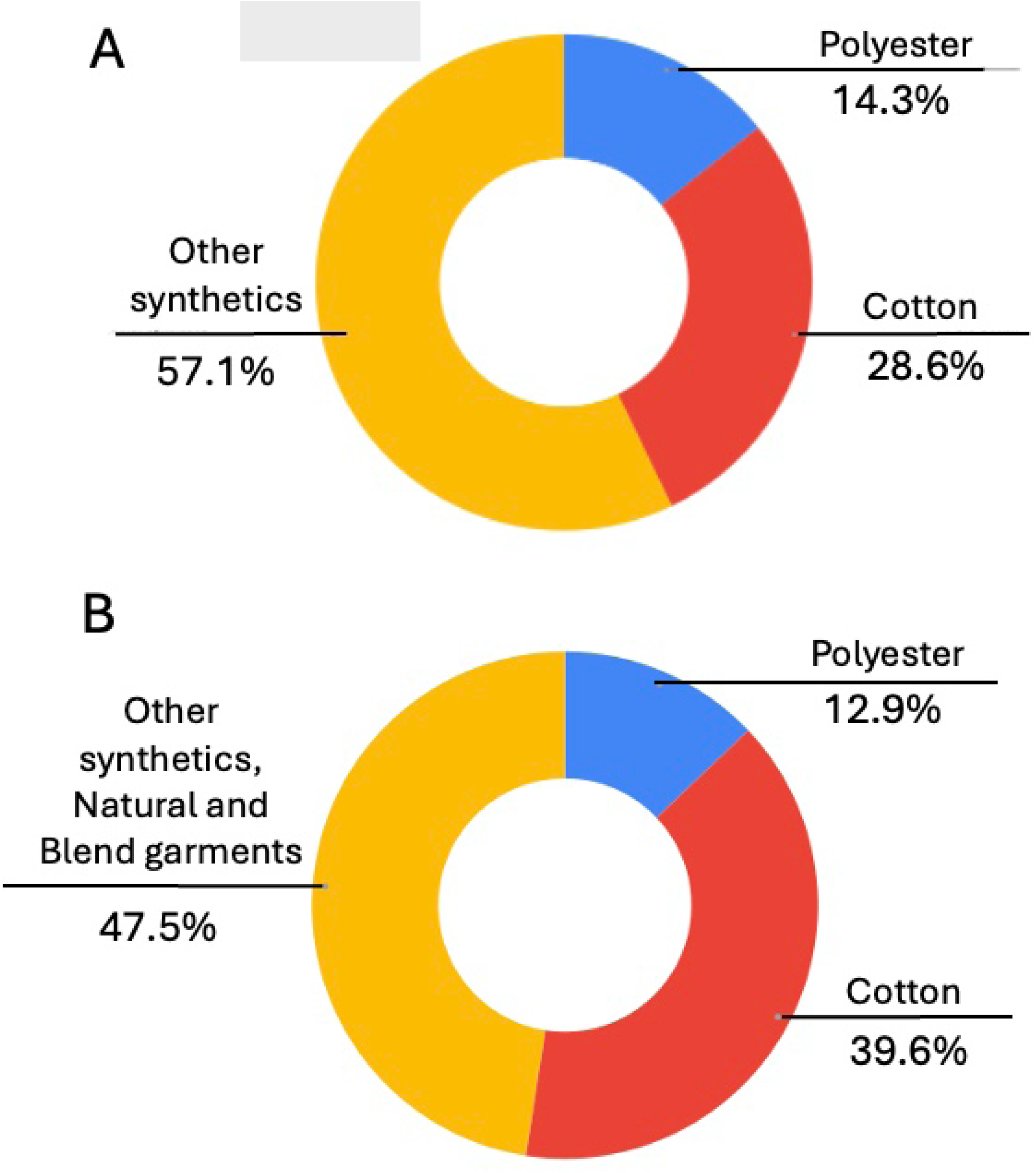
Pie chart representation of the percent occurrence of polymer types found in the microparticles collected in the lagoon water adjacent to the landfill (OKS 12.5, OKS20) (A, from InfraRed fingerprint analysis of microplastics) and those found in the waste pile of garments at Kantamanto Market (B, from material composition tags of garments).

*Textile waste showed similar brand assemblages between beach and market*. Collected tags showed that the 10 most common brands found in the textile waste were Marks & Spencer, Associated British Foods (Primark), Asda (George), Sainsbury’s (Tu), Next, Boohoo Group, Tesco (F&F), H&M Group, New Look Group, and GAP, representing up to 63% and 77% of the items found at the beach and in the market, respectively (Fig. 7). There was a 59% correlation of brands occurring between the two sites of waste collection, based on the tags analyzed, which was statistically significant (P < 0.05; Pearson ρ correlation coefficient).

**Figure 7.**
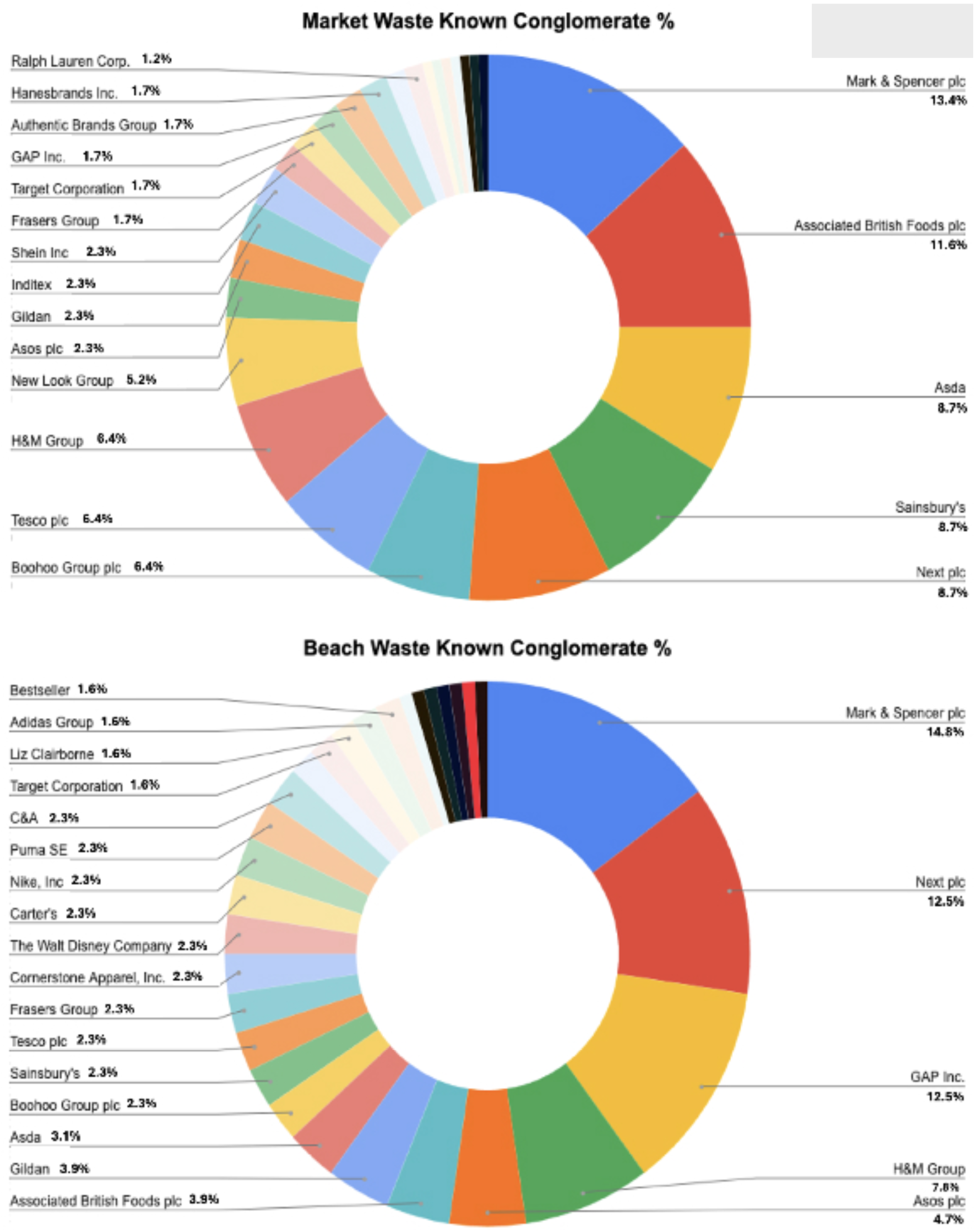
Pie chart representation of the percent occurrence of name tags for each brand found in the waste pile at Kantamanto Market (top) and found entangled and buried in the sand at the nearby beach (bottom). More than 60% of the tags originated from just 10 brands (Marks & Spencer, Associated British Foods, Asda, Sainsbury’s, Next, Boohoo Group, Tesco, H&M Group, New Look Group, and GAP).

## DISCUSSION

Our research around Kantamanto Market showed the extensive presence of microplastics and synthetic microfibers, in both the surrounding air and water, with concentrations seemingly representative of the varying attributes of the market activity. The data indicate that the concentrations of airborne microparticles we found varied depending on the day of the week and activities taking place on those days, with concentrations notably being higher after every Thursday, when bales of globally sourced secondhand clothing are delivered to the market. The concentrations were also higher during early morning on Saturdays, a busy market day when bales are opened and garments are displayed for sale in Kantamanto Market (Skinner 2019). Our study focused on microfibers, especially their synthetic version, because those are directly related to the secondhand textile activities in the market, yet keeping in mind that such microfibers could also have other origins (from outside the market or even a far-away location), while not dismissing the microplastic components, which can come from a diversity of sources (from tire wear to microplastics from burning waste, including textiles).

Our study showed, at least during the daytime when analyses were conducted, that microfibers that are released can float around and be carried to higher elevations (we tested at 10 m height), likely due to warm upward convection currents caused by heat generated by the hot surfaces heated by the sun, and to which the high number of people in the market can also contribute (Belzagui et al. 2019; De Falco et al. 2020; Periyasamy and Tehrani-Bagha 2022). Such upward hot air convection currents are known to occur in nature, where they help with plant seed dispersal for example (Inatsu et al. 2021; Mun et al. 2024) although they can also be very important in urban areas where they are considered by city managers for airflow improvements (Xie et al. 2006; Xue and Li 2017; Di Bernardino et al. 2018; Muniz-Gä<u>a</u>l et al. 2020). Here, we evidenced that the microfibers could be carried at least as high as 10 m into the air, although one cannot exclude that some of them could originate from other sources nearby as well. From there, they could potentially spread around with winds (we indeed found unweathered microfibers in neighborhoods around the market; see Online Resource 5) and fall back onto the ground at night, either when heat-driven convection currents stop or when they become entrained down onto the ground by rain events (Miao et al. 2023; Liu et al. 2025). Indeed, rainwater was polluted with microfibers (and microplastics), which was especially true for rain events occurring after 3-4 day periods of no rain. In such a case, we showed that the first flush of the rain (within the first hour or so, varying based on the rain intensity) could bring down microfibers in high concentrations. Practically, from a public health perspective, this indicates that people who collect rainwater for drinking usage should skip the first flush of the rain event, especially if it has not rained in several days, or should deploy filtration technology if any is available (like the clay filters commonly found across Africa).

Our data thus indicates that microplastics and microfibers can enter Korle Lagoon waters via two routes: One concentrated around the piles of discarded garments that shed fragments upon mechanical abrasion and weathering in the environment; and the other one more diffuse and associated with aerial deposition of airborne microplastics and microfibers, that can also be precipitated during rain events, and reach the lagoon via runoff and surface water drain outflows.

Ultimately, using Kantamanto as a case study representing the communities that receive, sort and process global fashion waste, we show that the high volume and low quality of garments arriving in these communities and the lack of infrastructure and capital to safely and efficiently deal with this waste, results in significant microfiber pollution. This pollution has the capacity to impact local air quality as well as local aquatic ecosystems and their associated services, thus potentially contributing to chronic public health issues (Manisalidis et al. 2020; Sridharan et al. 2021; Zhao et al. 2023; Song et al. 2024; Vasse and Melgert 2024).

*Global textiles as a source of airborne microparticles*. Using the PurpleAir sensors, we demonstrated that the mass concentrations of PM2.5 and PM10 particles were high and varied extensively around and near the market, decreasing when going further away from it. This indicated that the market was a source for airborne microparticles.

PM2.5 airborne microparticle mass concentrations in different neighborhoods in Accra have been reported to often be above the healthy air quality guidelines recommended by the World Health Organization, which is set at 10 μg/m^3^ (Zhou et al. 2011; Ofosu et al. 2012), while rarely surpassing 100 μg/m^3^ (Alli et al. 2021). Our study showed PM2.5 mass concentrations within the market can often reach levels up to between 125 and 175 μg/m^3^ (Online Resource 4A), sometimes peaking at much higher concentration levels around 1,300 μg/m^3^ (Fig. 2), which thus can be 10x-20x greater than for various areas in Accra. Because our study took place in the core of a textile market, and knowing that textiles shed microfibers (and/or fiber fragments that could be shorter and more like “elongated” microplastics) that can be <10 μm in size, it is fair to assume that our PM2.5 as well as PM10 mass concentrations could also include microfiber and/or shorter fiber fragments. This would be in addition to other microparticles typically found in cities (related for example to energy combustion, industry and road traffic) but also pollen and sand dust, often carried by sub-Saharan winds to Accra (Amegah et al. 2017).

Aside from the abnormally high airborne microparticle mass concentrations in Kantamanto, another peculiarity was that the PM10 mass concentrations were only about 1.0-1.2x greater than those of PM2.5. This was unlike other studies in Accra that showed PM10 values being 2.0-2.5x greater than PM2.5 (Arku et al. 2008; Dionisio et al. 2010). Because our PM10 mass concentrations were relatively elevated, the lower PM2.5/PM10 ratio in our study indicates that the source of PM2.5 is likely greater in the market environment, relative to its surroundings. PM2.5 values tend to be associated with byproducts of combustion such as those from fuel-powered engines (Arku et al. 2008; Dionisio et al. 2010). However, because the interior of the market (where measurements were made) is largely closed to vehicular traffic, the increased PM2.5 count concentration cannot be from on-site fuel-powered engines but likely stems from incomplete fire combustion of plastic material. Burning plastic waste is indeed a common practice in Kantamanto, which is done without wood, and not for the need of cooking or getting warmer, but for the need to discard waste and gain space. Such frequent (almost every night) and uncontrolled incineration of waste material is also done using textile waste, which is mostly made of plastic that, when burned, releases large numbers of PM2.5 microparticles (Bardales Cruz et al. 2022; Islam et al. 2022). Although another source of PM2.5 could originate from the individual use of cooking stoves across the market (Zhou et al. 2011), the importance of this practice is definitely less than the textile waste burning. The latter is likely the most prevalent indeed considering the many small fluorescent particles that were observed under epifluorescence in our study, which makes them synthetic (plastic), since black carbon microparticles from engine combustion and incineration practices with wood (both important sources of PM2.5 and PM10 microparticles in general) have not been previously reported to show native epifluorescence properties. In fact, additional staining methods are needed to image black carbon microparticles in fluorescence (Chen et al. 2019). The native fluorescence of microparticles in our study then indicates their synthetic (plastic) nature.

As expected, air quality in Accra shows daily variation relative to people’s activities, whether from cooking or driving to/from work (Alli et al. 2021), and/or from textile market activities and the burning of garments, as indicated by this study. We also showed here that airborne PM2.5 and PM10 microparticle concentrations decreased sharply in events of rain indicating that rainfall can be a major process to help microparticles, including microplastics and microfibers, transition from the airborne phase to the waterborne phase. This indicates that rain can be a major process by which airborne microfibers enter the waterways, in addition to wind driven processes that can deposit them onto water surfaces. Such an effect of rain bringing down airborne microplastics was recently described in other industrial settings (Do et al. 2023).

*Global textile waste as a source of microparticles for waterways*. We evidenced that Korle Lagoon is a receptacle of textile microfibers from runoff. Indeed, water samples collected from the lagoon showed high concentrations of microfibers (and other microplastics as well), with a spatial distribution along the lagoon and chemical makeup indicative of an origin associated with market activities. In particular, runoff following rain events and/or weathering fragmentation of garments discarded directly in the environment are likely a sustained source of microparticles for the lagoon. This is in addition to the microfibers probably reaching the water surface from airborne deposition. A majority of garments in the market contain polyester and/or cotton, which was also reflected in the waterborne microparticles, especially in waterways next to, and downstream from, the market. Our study thus highlights the extent to which the increase in textile production, and therefore textile waste, contributes to the release of microplastics and microfibers in the environment, either atmospheric or aquatic. This information is critical because microplastics and plastic microfibers are not biodegradable in a biologically relevant timeframe, making them accumulate in the environment where their long residence time makes them available to enter the food web and accumulate in organisms, including humans (Sharma et al. 2022; Thompson et al. 2024; Zhao et al. 2024). This, in turn, can affect the overall ecosystem services (Santana et al. 2017, Olmo-Gilabert et al. 2024).

*Concentrations of microfibers and microplastics were greater than in other parts of the world*. The concentrations of microplastics, including microfibers, found in our study, whether in the air or in the water, were usually by far the highest ever reported in an environment (e.g., Sazli et al. 2023). This does not come as a surprise considering that methodologies used can lead to underestimation of microplastics, both from airborne and waterborne sources (Xu et al. 2020). In addition, a direct comparison across studies is challenging because the methodology for sample collection and/or for microparticle counting usually varies from study to study, based on local environmental limitations, local expertise, the distinction (or not) between microplastics and microfibers, but also on the scientific question driving a particular study. Zhao et al. (2023) provided a relevant range of comparison for levels of airborne microplastics in megacities, with concentrations of microplastics possibly ranging broadly from <1 to >1,000 per m^3^, depending on environmental factors (e.g., winds, wildfires, smog conditions) and methods of collection (e.g., dry or wet passive atmospheric fallout vs. active suspended particle measurement). However, under normal circumstances and using suspended particle measurement as a reference (thus not brought down by rain or snow), others have measured relatively low concentrations, e.g. around 4 microplastics per m^3^ in Shanghai (Sridharan et al. 2021), 1.5 per m^3^ in Paris, and 0.6 m^3^ in the French Alps (Xu et al. 2024), often with no quantified distinction between microfiber and microplastic particles, except for Xu and colleagues who identified that 10% to 25% of the microplastics were microfibers. In our study, the Coriolis data showed about 100 to 700 microplastics per m^3^, thus about 100x what was found in Shanghai, and about 3 to 30 microfibers per m^3^ (considering only UV synthetic MFs, and not the BF MFs; see Online Resource 5), which is about 20x greater than in Shanghai and about 125x the numbers from the French Alps.

A similar analysis can be done with waterborne microfibers and microplastics. Microplastics in Kantamanto can originate from degrading plastic bags and disposable individual drinking water pouches (the main source of drinking water around the market), used extensively and discarded on the ground, with limited direct contribution from tire wear as otherwise found elsewhere in the city (Arku et al. 2008; Zhou et al. 2011; Knight et al. 2020; Amegah et al. 2022; Safo-Adu et al. 2024). As for microfibers their main source would be shedding from clothing (Carr 2017; Carney Almroth et al. 2018; Liu et al. 2019; Nowack et al. 2021; Liu et al. 2023). Concentrations of microfibers have been reported at around 2-30 counts per L in coastal waters (Suaria et al. 2020; Tang et al. 2023), which does not include lagoons as these have been much less investigated around the world for their content in microfibers. The same can be said for microplastics, although in this case one comprehensive study gathered data from the literature since 2000 (Garcés-Ordóñez et al. 2022), showing concentrations of 20 to 90 counts per L on average for lagoons around the world, with about 40% of these numbers attributed to microfibers, thus making the count to about 12-54 microplastics per L and thus about 8-36 microfibers per L. This is in line with studies reporting up to 30 counts of microplastics per L in harbors and with 0.5-2.0 counts per L in coastal and more open ocean waters (Suaria et al. 2020; Tsang et al. 2020). The difference in concentrations relative to what we found in Korle Lagoon waters (between 100 to 1,600 microfibers per L and between 1,000 to 11,000 microplastics per L; Online Resource 8) is staggering and likely due to the source being the abundance of textiles in the market as well as in the dumpsite (as waste) along the lagoon. Such high concentrations (up to 25,000 counts per L) were previously observed in lagoons in the Florida Keys (USA) (Badylak et al. 2021) and were considered the result of debris (mostly polystyrene pellets) coming from recent hurricane activities and accumulation in specific areas of lagoons due to winds and currents. This was not the case for the data collected in Korle Lagoon, which shows a slow but steady flow to the ocean.

Another factor possibly contributing to the difference in concentrations of microplastics found across studies could also originate from the differences in methods used to collect water samples and to analyze the water samples. Indeed, a common way to collect microplastics involves a Manta net, which is not easily used in a shallow lagoon (Royer and Deheyn 2019); this method has also been documented to underestimate the numbers of microfibers that can pass through the mesh due to their elongated shape, and because the mesh size is usually larger, around 500 μm (Barrows et al. 2018; Du et al. 2022; Concato et al. 2023). Hence, the technique to collect set volumes of water for subsequent filtration and counts of microfibers (and microplastics) retained on filters, is likely more representative of the actual waterborne microfiber concentrations.

*Textile waste from global sources passing through Kantamanto Market contaminates the coastal ecosystem.* Beaches in Accra have masses of entangled garments that arise on and under the sand. Composed of dozens of garments, these masses develop into long tentacles (bundles as much as 30 cm in diameter and meters long) that go deep into the sand, and their extraction, through beach cleanups organized weekly by The Or Foundation, represent a monumental effort. It is unclear how long it takes for garments to form tentacles like these with the back and forth of the waves and tides, and for them to be buried deep into the beach. However, the local coastal community, including The Or Foundation’s Beach Monitoring team, which counts the tentacles every week, has observed them for several years (The Or Foundation 2022). Taking tags from garments on the beach and comparing them to the tags from the textile waste in the market demonstrated that garments found at the beach originate as part of the global secondhand clothing trade passing through Kantamanto.

The tags showed that the 10 most abundant brands found at the beach and as market waste correlate well with one another. While it is possible that some of the garments are trashed directly at the beach after purchase and use for a little while, it is unlikely that such “isolated” circumstances would lead to the formation of the long tentacles that have been observed.

Brands of the garments most commonly found as waste on the beach and as waste within Kantamanto Market were from the Global North, with companies headquartered and distributed principally within the USA, the EU and the UK. Most of these companies are well known to be involved in the fast fashion economy (Boohoo, Gap, George, H&M, New Look, Next, Primark), although not always recognized as such by consumers (as is the case for Marks & Spencer, for example). Thus, within the fast fashion economy, whereby consumers are influenced to rapidly change wardrobes at relatively affordable prices (The Economist 2015), garments can be considered environmental pollutants of emerging concern when discarded as waste at their end of life due to the rapid turnover and high volumes of garments in circulation.

*A simple collection and processing technique to enable citizen science*. In this study, we worked together with the local community to access sites for collecting samples, to engage community leaders to implement new waste management practices, and to empower citizens to help with sample collection. As such, we used a simple and straightforward technique that easily could be performed by any citizen, with only brief training needed. The technique is effective in collecting and quantifying microfibers, but also microplastics. The technique relies on small amounts of water (50 mL) that can easily be sampled by scientists and citizen scientists alike using standardized tubes, making this attractive for large sampling collection involving the community. The use of water volumes greater than 50 mL was tested as well (including 1,000 mL, 500 mL, 200 mL; data not shown) but shown to be more challenging due simply to the larger volume to handle (with increased risks of clogging). It was also more challenging for counting the microfibers, due to excess debris on the filters, or due to difficulty counting because of the excessive number of microfibers overlapping each other in the field of view. It must be noted that in some instances here, the 50 mL volume had to be analyzed 10 mL at a time, using several filters, to facilitate accurate counting and then added together at the end to represent the one sample (see Online Resources for more details). The proposed technique thus provides flexibility when dealing with water masses that are heavily charged with silt, plankton and/or debris, which can clog the filtration process early on (before reaching the complete 50 mL volume). The proposed technique (which by default uses triplicates of 50 mL volumes) thus complements the most common one used for microplastics in particular, using Manta Net towing/trawling, which is bulky, requires a boat, and is thus difficult for citizens to access.

## CONCLUSION

This citizen science-led study provides a comprehensive assessment of microplastic and microfiber concentrations in the air, rainwater and lagoon water around Kantamanto Market and its known dumpsites of global secondhand textile waste. The global textiles and textile waste passing through the market appear to be a significant source of microfibers (and microplastics), with concentrations within and around the market sometimes more than 100x greater than in other metropolises around the world. We demonstrated the dynamic distribution of microparticles following their generation in the market, with particles in the air precipitating back down with rain events, and possibly with wind and cooler temperatures. The effects of global textile waste passing through the market and entering the surrounding environment go beyond the microscopic particles. Indeed, we showed a strong correlation between garments considered waste at the market and the ones found on the surrounding beaches, by comparing brand composition of the garments. Overall, we emphasize the fact that fast fashion contributes to poor water and air quality, which is likely the case across other densely populated cities worldwide considering that the processes we highlighted in Kantamanto are not site specific, and could take place in other secondhand clothing markets. As such, the study highlights the global need for new policy regulations and programs to ensure that cleanup programs and proper waste management efforts are sufficiently funded along the entire life cycle of the garments, including for existing and future waste, repurposing and recycling streams.

## GLOSSARY

Fast fashion: An approach to the design, creation, and marketing of clothing fashions that emphasizes making fashion trends quickly and cheaply available to consumers. Fast fashion is characterized by overproduction and the use of cheap (often exclusively synthetic) materials and fabrication techniques.
Fiber fragment: A microscopic fragment that arises from the shedding or degradation of fiber threads from either natural or plastic (synthetic) textile materials. Fiber fragments are characterized by a diameter usually around 8-12 μm and a length that can be greater than the diameter, in which case it would be known as “microfiber.” Shorter fiber fragments (with diameter shorter than their length) would then be reported as microplastics.
Kantamanto Market: The world’s largest secondhand clothing and upcyling market, located in Accra, Ghana, and featuring over 10,000 vendor stalls.
Microfiber: A microscopic fragment that arises from the shedding or disintegration (physical break off) of both natural and plastic textile thread materials. Also known as a “fiber fragment.”
Microparticle: Any microscopic particle, regardless of shape, whether synthetic or originating directly from nature (mineral dust or small organic debris).
Microplastic: A microscopic fragment that arises from the disintegration of plastic materials, or that is made small to begin with, such as microplastic beads from cosmetics. As used here, the term “microplastic” is distinct from “microfiber”, although microplastics could be short microfibers (see fiber fragment definition above) As such, in this study, microplastic counts represent something distinct from microfiber counts.
Natural microfiber: A microscopic fiber fragment shed from a natural textile material, such as cotton, wool, or other plant-based sources.
Plastic (or synthetic) microfiber: A type of microplastic with a “fiber shape”. Plastic microfibers typically arise from the shedding or disintegration of textiles, although they can also originate from tire wear, carpets or upholstery.

## Author’s contribution

DDD and BS conceptualized the project. JA performed the field and laboratory work in Ghana, under DDD guidance. DDD performed analysis in California. DDD wrote the first draft of the paper, which was then reviewed and finalized by BS and JA, and several members of The Or Foundation.

The authors declare that they have no known competing financial interests or personal relationships that could have appeared to influence the work reported in this paper.

## Authors biography

Dimitri Deheyn, PhD

Dimitri is a Research Scientist at the Marine Biology Research Division of Scripps Institution of Oceanography, University of California San Diego, where he conducts cross-disciplinary research on biomimicry. In this frame, he studies alternative materials to emerging pollutants, which includes plastics, textile microfibers and associated additives. Some key steps of his studies include assessing plastic and fiber biodegradation (from both natural and synthetic fibers, untreated and treated with additives). His research has a global perspective, across a diversity of ecosystems, around the world. Dimitri grew up in Africa, and earned his PhD in marine sciences and ecotoxicology from the Free-Thinking University in Brussels, Belgium where he started his academic career.

Joe Ayesu

Joe is an integral member of the Accra Arts Centre beachfront community, where he has a shop selling works from across West Africa’s arts and crafts industry, including many of his own creations. Living and working in this location for decades he has observed Accra’s waste management challenges firsthand, particularly those related to textiles. Joe majored in science in his secondary school days and studied the fundamentals of medical laboratory technology at Accra Technical University. He manages The Or Foundation’s team of citizen scientists, mobilizing dozens of people for beach monitoring and cleanup efforts, sample collection and imaging.

The Or Foundation

Headquartered in Accra, Ghana, The Or Foundation is the leading not-for-profit implementing Globally Accountable Extended Producer Responsibility for textiles through a voluntary program in service of Accra’s Kantamanto Market community, the largest secondhand clothing market in the world. Working at the intersection of environmental justice, education and fashion innovation since 2011, the mission of the NGO is to identify and manifest alternatives to the dominant model of fashion – alternatives that bring forth ecological prosperity, as opposed to destruction, and that inspire citizens to form a relationship with fashion that extends beyond their role as consumer.

Branson Skinner

Branson was studying in Ghana when he was confronted with the impact of textile waste on the environment and communities in the country. This led him to co-found The Or Foundation, a Ghana-based non-profit working at the intersection of environmental justice, education and fashion development. At The Or Foundation, Branson has combined his diverse background as an educator, farmer, urban planner, and musician to work closely with the local community in Accra and lead the organization’s work implementing innovative solutions and science-backed research initiatives to address fashion’s global waste crisis. Branson studied Globalization of Public Health and Societies at NYU Gallatin, where he was a Reynolds Scholar for Social Entrepreneurship. He also holds a Masters of Community Planning degree from the University of Cincinnati.

## Acknowledgments

This work benefited from the work of some volunteers at Scripps Oceanography, including Layla Nazif and Avery Dougherty, to whom we are thankful for their time. In Ghana the work has been greatly facilitated by the Accra Metropolitan Assembly, the Kantamanto Used Clothing Sellers Association, the Kantamanto Hard Workers Association, and the ADAMOG group in Old Fadama. Funding for the project was provided to BS from The Or Foundation, leveraging financial support from the SHEIN Global EPR Fund; The Biomimicry Institute; Soliii Earth; and general public contributions.

## Notes

### Competing Interest Statement

The authors have declared no competing interest.

### Summary of Updates

This paper has been revised mainly around the text, by substituting numbers of particles by concentrations. Changes throughout the text also provide more details on the methods (with an extensive set of data added to the supplementary information) and also provide more justification about some of the methods we used.

