## Supplementary material for "Microplastics found in high concentrations around Kantamanto, the world’s largest secondhand textile market in Accra, Ghana: A citizen science study": Supplemetary information

♦ Contributed equally to the work.

### MATERIAL AND METHODS

Sample processing for microplastics and microfibers concentrations followed a routine procedure from the Deheyn Lab, which has been put to practice over the last few years from samples from all over the world (New York Times 2020).

*Water sample preparation*. In the laboratory of The Or Foundation (in Accra, Ghana), the content of the Falcon tube was vortexed, and the water-containing-particles were transferred to a 50 mL syringe (the exact volume was identified then) for filtration on a Advantec GF/F FiberGlass filter of 2.5 cm in diameter and 0.6 μm pore size. Although in most cases the entire 50 mL of the sample could be filtered with gentle pressure on the syringe, in some cases the filter would rapidly start clogging after just past a few mL (the water could be very turbid). The filtration process was then stopped to avoid cracking of the filter and forcing water and material through, and the filter replaced with a new one. The process was repeated as many times as needed for the entire volume (50 mL) to be filtered in as many filters as needed. The numbers of items per filter were then added to one another to represent the entire sample. In some situations, especially for rainwater, the water volume was less than 50 mL. In this case, the filtered volume was recorded, and the number of items counted on the filter expressed relative to 50 mL. As such, all numbers of microparticles we reported were for the same volume unit, which allowed for comparison across samples. Thus for water samples, the concentrations across this type of unique sample collection and process were always expressed as a concentration of counts/50 mL.

Each filter was then placed on a small piece of aluminum foil that was folded back onto the filter without touching it, shielding the filter from direct deposition of airborne contamination, and to avoid losing material by direct contact with the foil. The filters were let sit in a drawer, overnight at least until dry (the filter is then stiff and can easily be moved around using metal forceps). See details in Online Resource 2, below.

This process was done for 3 of the 4 replicates as the 4$^{th}$ replicate was kept in the fridge to be sent later to the Deheyn Lab (Scripps Oceanography, CA), where the same process was conducted, which served as an independent quality control. Data showed similar results between the two laboratories.

*Microparticle counting from filters.* All counts of microplastics and microfibers gathered from each of the digital images of the filters were made by at least 4 trained individuals. We used the brightfield digital images to count colored items, used here as a proxy for their young and unweathered stage, thus indicative that they were recently released in the environment. We used the fluorescent digital images to count synthetic microfibers and microplastics, which most fluoresce. All microfibers (elongated items) were counted. As for microplastics (particulate items), they were all counted as well, unless found in overwhelming numbers spread homogeneously across the filters. In such case, the counts were made from a fourth of the filter, and the numbers multiplied accordingly for full filter count.

All items found with red fluorescence were by default considered natural microfibers or microparticles (the red fluorescence originating from chlorophyll), unless it was obvious from the corresponding brightfield images that these items showed colored dyes. In this case, these items were categorized as synthetic material. See details in Online Resource 1, below.

Overall, the data showed that the contribution of freshly released microparticles (bright field counts) was low compared to their brightly fluorescent analogs (epifluorescent counts).

*Polymer identification of microparticles*. Sample processing for chemical identification of the polymer types was performed on the most abundant microparticles, especially the ones that clearly showed a fiber-shape since we wanted to focus on textile materials most importantly. The polymer chemical identification was done by Quantum Cascade Laser InfraRed (QCL-IR) spectral imaging, using the Spero® IR system (DLS DayLightSolutions Inc.). This technique is well suited for the purpose of microplastics chemical identification (Primpke et al. 2020).

For such analysis, filters showing high and/or large numbers of microparticles were placed back under the SMZ 1800 stereoscope and observed using a simultaneous combination of bright light and epifluorescence (as used for imaging to get microparticles counts). With ultrathin high-precision forceps, this illumination setting allowed the observer to hand pick microparticles that were either colored or not, then possibly emitting fluorescence. A total of 35 microparticles (some were clearly microfibers while some were longer than wider, thus considered "elongated microplastics") were collected and placed on an IR coated Kevley slide for imaging under the Spero®.

The generated infrared hyperspectral cubes ranged from 950 to 1800 $cm^{-1}$, which is a typical IR fingerprint range of polymers. Spectra were selected from each object in the field of view by selecting an in-focus area, which usually did not exceed 15 μm/side. The generated spectrum from such area was processed and displayed using the DLS built-in ChemVision software. This process was performed in several areas of the object to ensure that the collected spectra were showing good signals (smooth spectral lines) and were comprise within the relative logarithmic intensity range of about 2-3 (a value >4 was considered saturated).

The collected spectra were then analyzed for their absorbance "valleys" which were compared to the various reference standards (Online Resource 3). These include typical plastic material used for packaging but also reference material most specifically tailored to the textile industry); all these reference standards were white. While the plastic reference standards were purchased (Polymer Kit 1.0), the textile references were provided by Lenzing Inc (https://www.lenzing.com/) and Archroma (https://www.archroma.com/) with whom Deheyn as ongoing collaboration. All the reference materials were processed using the Spero® in the Deheyn Lab, to ensure methodology of analysis was identical to that of the samples. The spectra from the samples were then compared for alignment of absorbance valleys in Excel with the spectra of the references for polymer identification (Online Resource 3).

*Textile waste by brand tag*. To assess the relationship between the textile waste (this time as full garments) from Kantamanto Market and the textile waste pollution found in the lagoon and at the beaches flanking the opening of the lagoon, we compared the brand tags from 100+ garments found at the market versus the beach.

By working with secondhand clothing retailers within Kantamanto Market, we know that full garments are indeed discarded at the market, ending in the lagoon waters, where they sink (and can tumble slowly midwater or on the seafloor without being stopped by the surface barrier in OKS20; see paper) or in adjacent storm drainage systems flowing out to the coast. Our hypothesis therefore is that the sunken textile waste progressively makes its way out of the lagoon and into the ocean, from where the waste is distributed along the beaches. The textile waste on the beach would therefore originate mainly from the Kantamanto Market, which should be reflected in similarities between the profile of tags found from the garments, considering the caveat of course, that the time for waste garments to travel from the market to the beach via the lagoon or other urban drainage systems remains unknown, and probably highly variable from item to item (e.g., due to different sizes, different shapes, different materials).

Textile waste by brand tags from the market was determined by a process that started each week, by speaking with around 10 market retailers (different each week). This was to inform the retailer about the study, and the willingness for them to keep for a week the items that could not be sold and that they were going to discard (as textile waste). We would then return to the retailers to acquire data from the textile waste, which we recorded on site directly through notes and photographs. Sorting through the textile waste, we recorded the garment type (e.g., Men's Shirts, Women's Joggers), the fiber type of each garment (cotton vs. polyester vs. nylon or other types, identified brand tag of each garment.

Textile waste by brand tags from the beach was determined by weekly beach cleanups organized by The Or Foundation. These involved digging into the sand to extract the numerous and heavy textile tentacles, defined here as entangled pieces of garments forming together long threads rooted deep into the sand. After the sand is rinsed off the textile tentacles in the ocean, team members would sift through the garments and remove brand tags. These were brought back to the lab, let dry, photographed and their abundance recorded as above. Using Excel, a Pearson correlation coefficient was then tested for significance between the numbers of brand tags collected from the waste pile of the market, and those found buried in the beach.

### Online Resource caption

**Online Resource 1.** Testing the fluorescence property of white and color dyed materials of synthetic origin relative to natural materials, including natural plant debris or phytoplankton samples. Left panel represents brightfield images with the corresponding images in epifluorescence on the right. When the epifluorescence signal was dim (not visible), the exposure time was increased 100x (insert image). All images were at 8x magnification except the PES fibers with leaf debris (0.75x magnification). All imaging conditions remained unchanged from the white PES imaging (except for the inserts)

We used reference fabric material (white and dyed) from Lenzing Inc and Archroma to assess the fluorescence capacity of the various polymers commonly used in textiles, which included natural polymers such as cotton, lyocell and modal, as well as synthetic polymers such polyester, nylon, elastane and cellulose acetate.

We found that the synthetic materials produced vivid blue/cyan blue fluorescence when colored in white or light hues (such as green or yellow) while their fluorescence is sharply reduced when colored with darker dyes (brown or black), then requiring 100x longer exposure times to detect any fluorescence signal (Online Resource 1A).

In contrast, none of the natural materials produced fluorescence under the same settings used for the white synthetic materials, with any fluorescence detected only when using 100x longer exposure times (Online Resource 1B).

Considering that we expected our samples to also contain debris of plants or algae, we exposed synthetic material (fibers/fragments) to samples of debris of crushed leaves of plants, and phytoplankton from cultures (available in the Deheyn lab). Under fluorescence, the synthetic fibers/fragments produced the expected cyan blue/blue fluorescence, while the plant and phytoplankton produced vivid red (Online Resource 1C).

Abbreviations: AC: Cellulose Acetate; CO: Cotton; CLY: Lyocell (wood-based); CMD (wood-based); EA: Elastane (spandex); PES: Polyester; PA: Nylon.

**Online Resource 2.** Example of images taken from filters used to retain microplastics and microfibers from samples. Imaging was performed from each of the dried filters. The imaging used a 0.75x objective to allow for the entire filter to be seen in the field of view and be quickly imaged (in less than a couple of minutes), which reduced exposure to possible airborne microplastics and microfibers contamination.

The imaging was performed in brightfield first and followed immediately by epifluorescence imaging (ex. 360-380 nm, em. >415 nm Long Pass filter) without the filter being moved. In Ghana, we used a Motic SMZ-171 stereomicroscope equipped with color digital CCD camera Exelis MPX-20C. In California, we used a Nikon SMZ1800 stereomicroscope equipped with color digital CCD Nikon camera. Images were comparable between the two settings.

All counts were made from each digital image, by at least 4 individuals (3 in Ghana and 1 in California USA). We used the brightfield digital images to count colored items, and the fluorescent digital images to count items with blue vs. red fluorescence. All microfibers were counted while for microplastics, they were all counted unless found in overwhelming numbers spread homogeneously across the filters. In such case, the counts were made from a fourth of the filter, and the numbers multiplied accordingly for full filter count. All counts were expressed by the volume of sample collected, namely 50mL, thus with concentration expressed as “counts/50 mL”.

The illumination conditions were set across all samples to best detect the brightest signal from items on the filters (thus to detect specific color in bright field mode versus to best see blue/cyan blue color in fluorescence mode).

As such, our imaging permitted to identify four categories of items from the images: (1) the ones colored and visible in bright field that represented recently released items from materials (since their color was not bleached yet by UV weathering) and that could possibly produce various shades of fluorescence, (2) the ones producing bright red fluorescence that we associated with being natural plant micro-debris, which could be confirmed from the bright field imaging as well, (3) blue/cyan blue fluorescence microparticles representing microplastics, and (4) blue/cyan blue fluorescence elongated items representing microfiber synthetic material.

One must note that the blue/cyan blue fluorescence was mainly associated with materials of white color, which might be uncolored to begin with or originally colored but then weathered and the color bleached. This made the material have different shades of whites, which came with different levels of fluorescence. This observation was not investigated further in this study.

Abbreviations: MFs: Microfibers. MPs: Microplastics.

**Online Resource 3**. Examples of infrared spectra from the QCL-IR Spero® analysis (using ChemVision for spectral display), showing the reference spectrum (black bold) and the spectrum for 3 different microparticle isolated from a filter. The subtle variations between the control spectrum and the "field" samples are representative of the unknow history of the field samples in terms of their age, weathering and original form.

Abbreviations: CO: Cotton; PEST: Polyester; PP: PolyPropylene.

**Online Resource 4.** Data from the PurpleAir® sensor. **A.** PM2.5 and PM10 particles mass concentration (box plots, in µg/$m^3$) across the various sites of investigation showing that the counts were systematically greater in the market, and, within the market, greater at 10 m compared to 5 m. **B.** PM2.5 and PM10 particle count concentrations (box plots, counts/L) showing the same trend as for the mass concentration, but with values of PM10 microparticles about 10x less concentrated than PM2.5 microparticles. The fact that PM10 microparticles including microfibers were found away from the market suggests that microfibers could be taken up by winds and carried away from the market. This was supported by the fact that other data from the Coriolis found microfibers (weathered and unweathered) off the market as well (see Online Resource 5). Data were collected every minute continuously on May 29, 2023. We took the Median values of 2 min consecutive segments, thus giving 30 numbers per hour, with a total of N=720 for each box plot.

Abbreviations: KM: Kantamanto Market, open area with passages for people between stalls. HQ: Headquarters of The Or Foundation just 1 km from the center of the market; OSU: Residential neighborhood in central Accra, about 5 km from the market.

**Online Resource 5.** Data from the Coriolis® sensor (6,000 L of air analyzed each time). Airborne synthetic microfiber concentrations (box plots; counts/6,000 L) across the various sites of investigation showing that the microparticle concentrations were greater in the market and within the market, greater at 10 m compare to 5 m. Data were collected with the Coriolis air sampler for 2 hrs each time. N=9, from 3 sample replicates each counted by 3 different individuals.

Abbreviations: HQ: Head Quarters of The Or Foundation; WH: WareHouse, large, enclosed space within the market and where textile waste is treated for reuse; KM: Kantamanto Market, open area with passage of people between stalls.

**Online Resource 6.** Rainwater synthetic microfiber concentrations (box plots, counts/50 mL) collected with a Stratus Rain Gauge (https://www.scientificsales.com/6330-Stratus-Rain-Gauge-p/6330.htm) set laterally 70 cm away from an ATMOS41 weather station on the roof of The Or Foundation Headquarter. Samples correspond to different rain events, each with their own volumes (X axis). The volume of the rainwater was measured in mL, and recorded in mm by the weather station. However, the weather station recorded data 24/7, while the rain gauge collected water during different time frames that were relevant and safe for the citizen scientists to set it out and bring it back in. Data counts are expressed per set volume of 50 mL since our focus was on the actual volume collected rather than the record of precipitation. N=9, from 3 sample replicates each counted by 3 different individuals. The concentration of synthetic microfibers was high for short rain events (within 2hr long) and decreased steadily as rainfall got heavier and/or lasted longer. *This rainwater was collected the same day as the 185 mL rain event, but just a few hours later. This shows that the concentration of synthetic microfibers in volume of rainwater depends on precedent episodes of rain.

**Online Resource 7.** Rainwater synthetic microfiber concentrations (box plots, counts/50 mL) for rain events in relation to the time (number of days) since the last rain event. Data show that the concentration of synthetic microfibers increases greatly in the rainwater, after 4 days with no rain. N=9, from 3 sample replicates each counted by 3 different individuals. Data counts are expressed per set volume of 50 mL.

**Online Resource 8.** Synthetic microfiber concentrations (box plots, counts/50 mL) from Korle lagoon water showing that site flanking directly the market in terms of runoff (OKS13) has the greatest counts, which increased even further at the site upstream from the surface barrier (OKS20) where floating debris and microparticles accumulated. N=9, from 3 sample replicates each counted by 3 different individuals. Data counts are expressed per set volume of 50 mL.

**Online Resource 9 (Table).** Minimum (Min), Maximum (Max), Mean and Median values for the counts (all from 50mL) of microfibers and microplastics identified as colored in bright field (freshly released) or with red fluorescence in epifluorescence (natural), from samples of air, rainwater, and Korle lagoon water. For all counts, N=9, except for the airborne sampling counts (N=3). Data counts are all expressed per set volume of 50 mL.

Abbreviations: HQ: HeadQuarter of The Or Foundation (roof) at two different times (a,b); WH: WareHouse space adjacent to the market at two different heights (5 m, 10 m); KM Kantamanto Market (center) at two different heights (5 m, 10 m). All lagoon sample counts originate from 50 mL of surface water.

Online Resource 1A

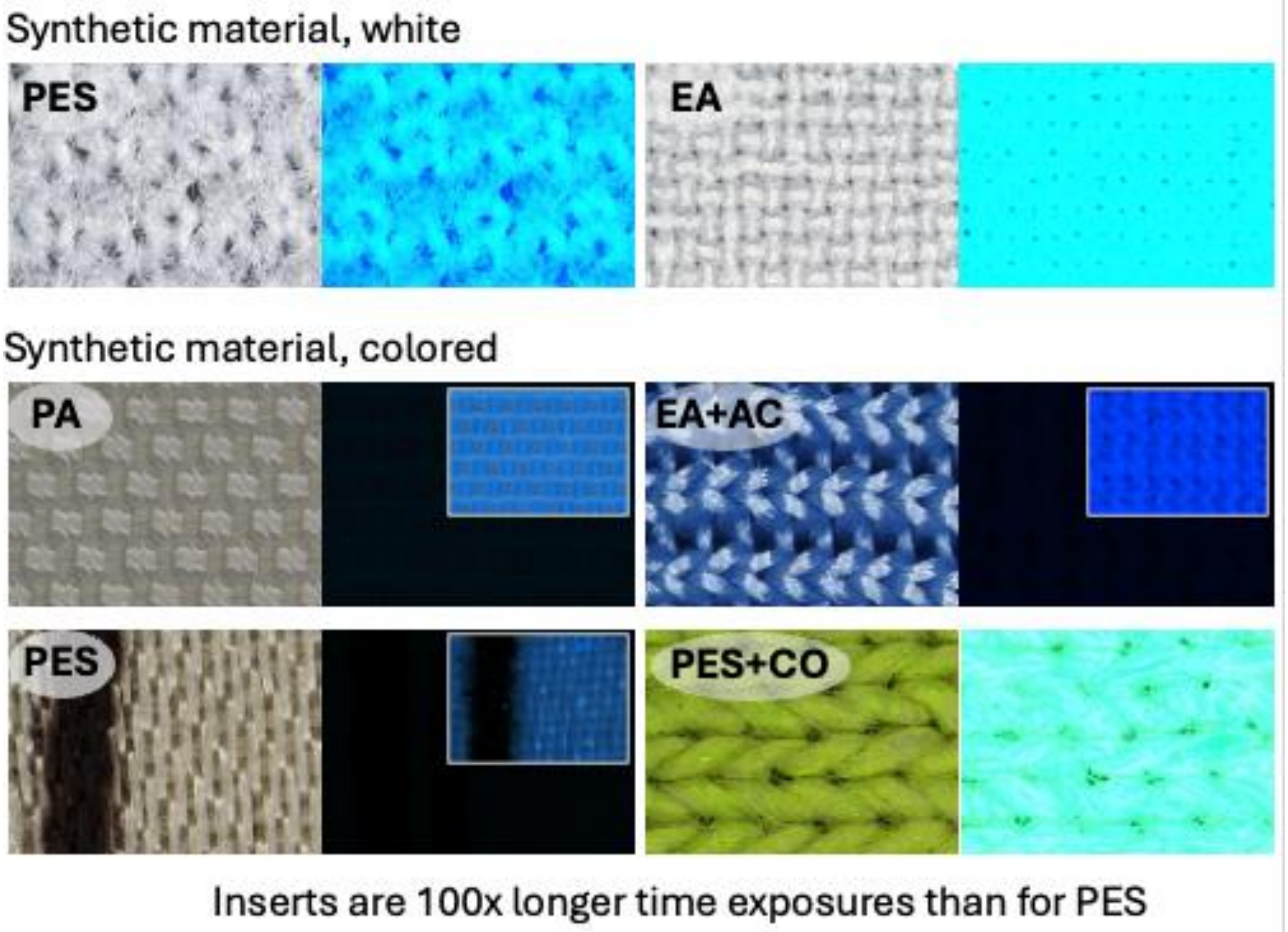

Online Resource 1B

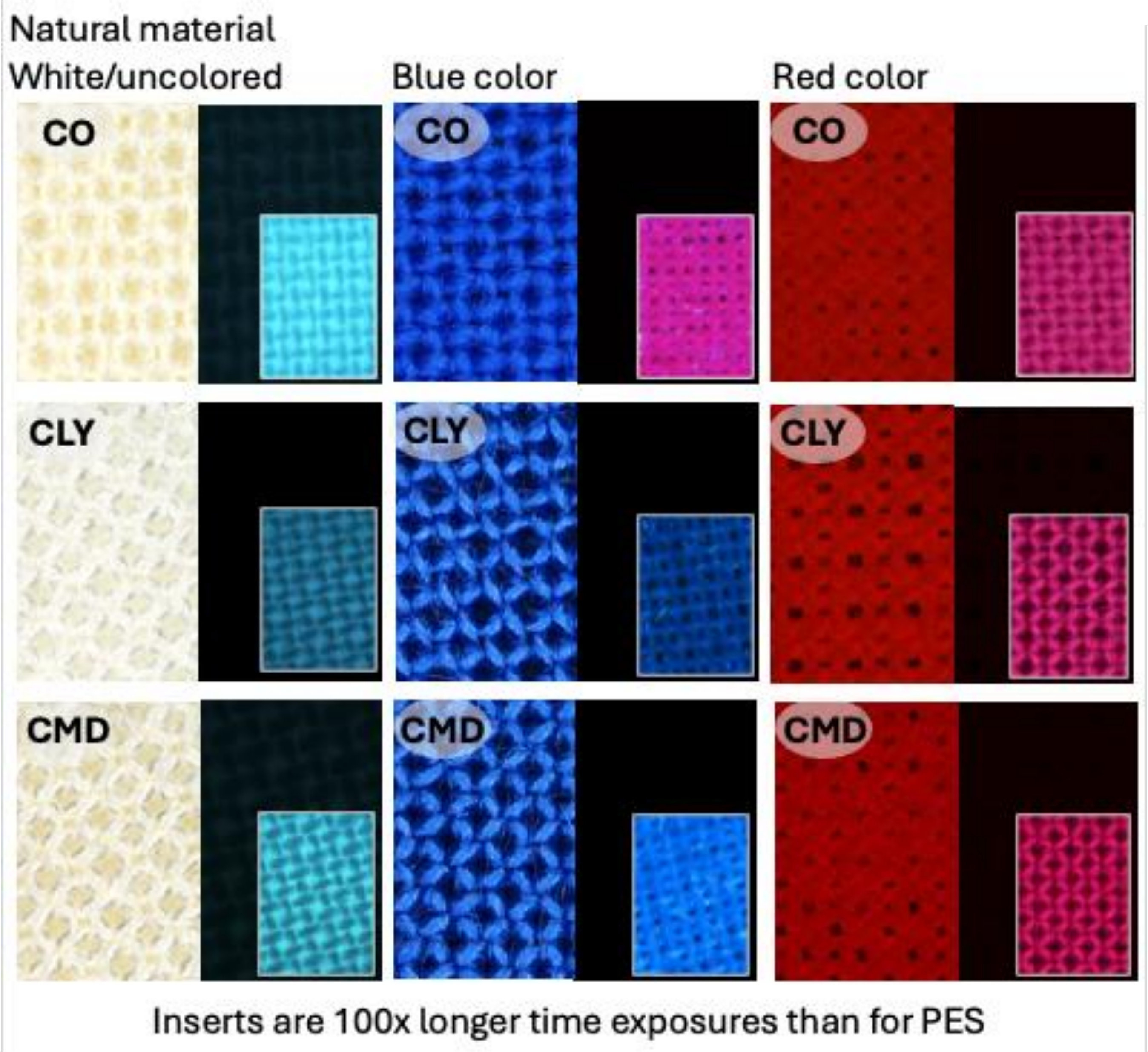

Online Resource 1C

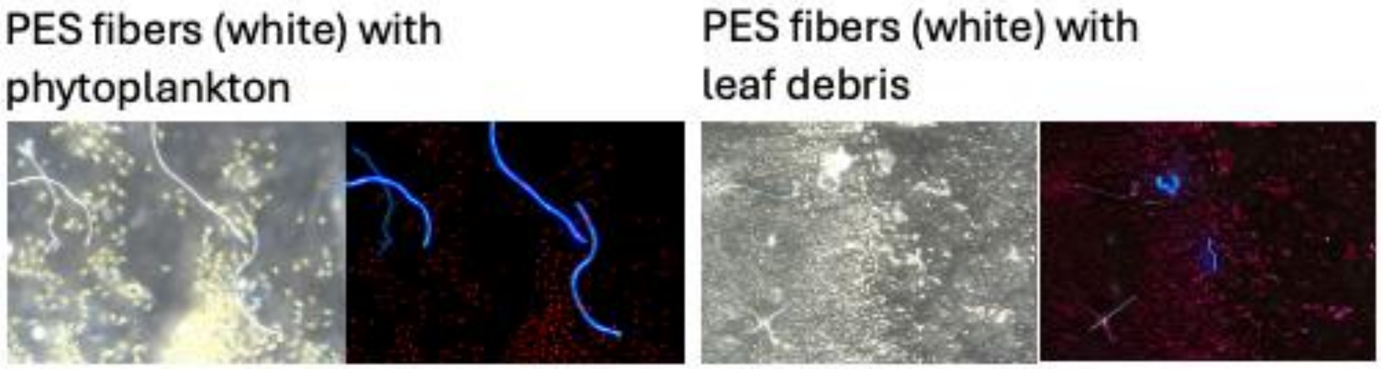

Online Resource 2

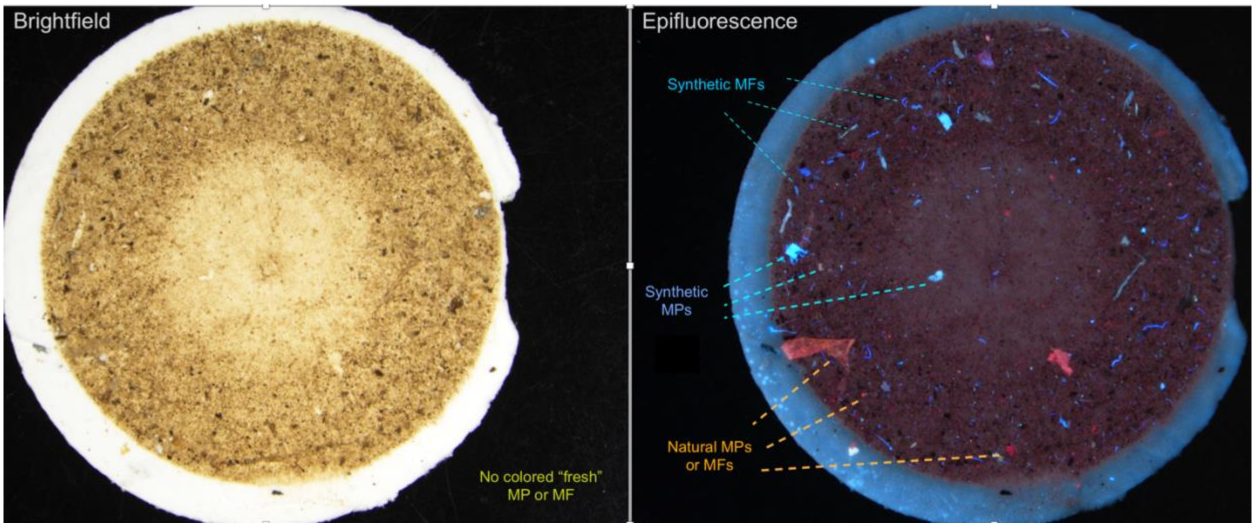

Online Resource 3

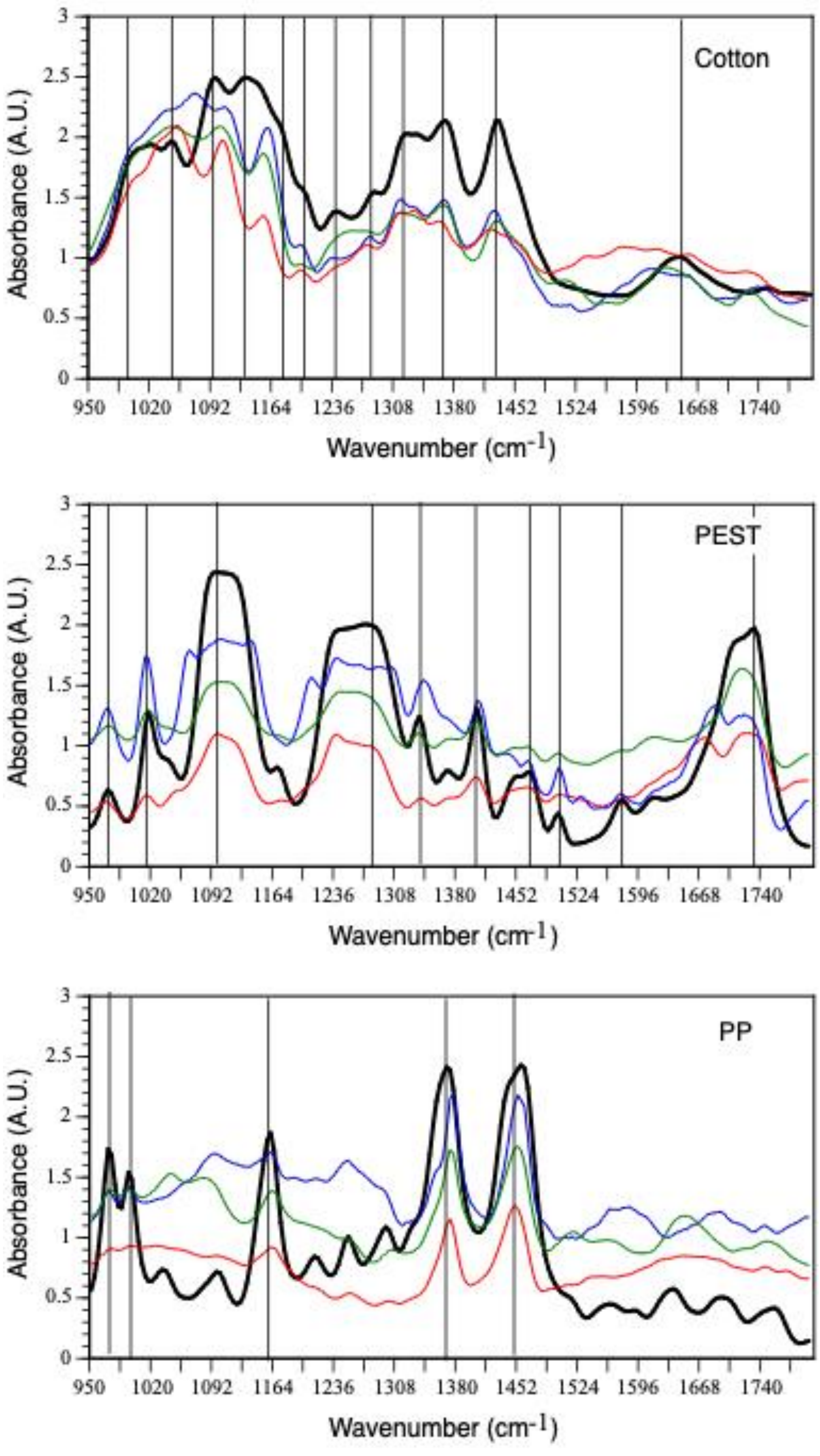

Online Resource 4

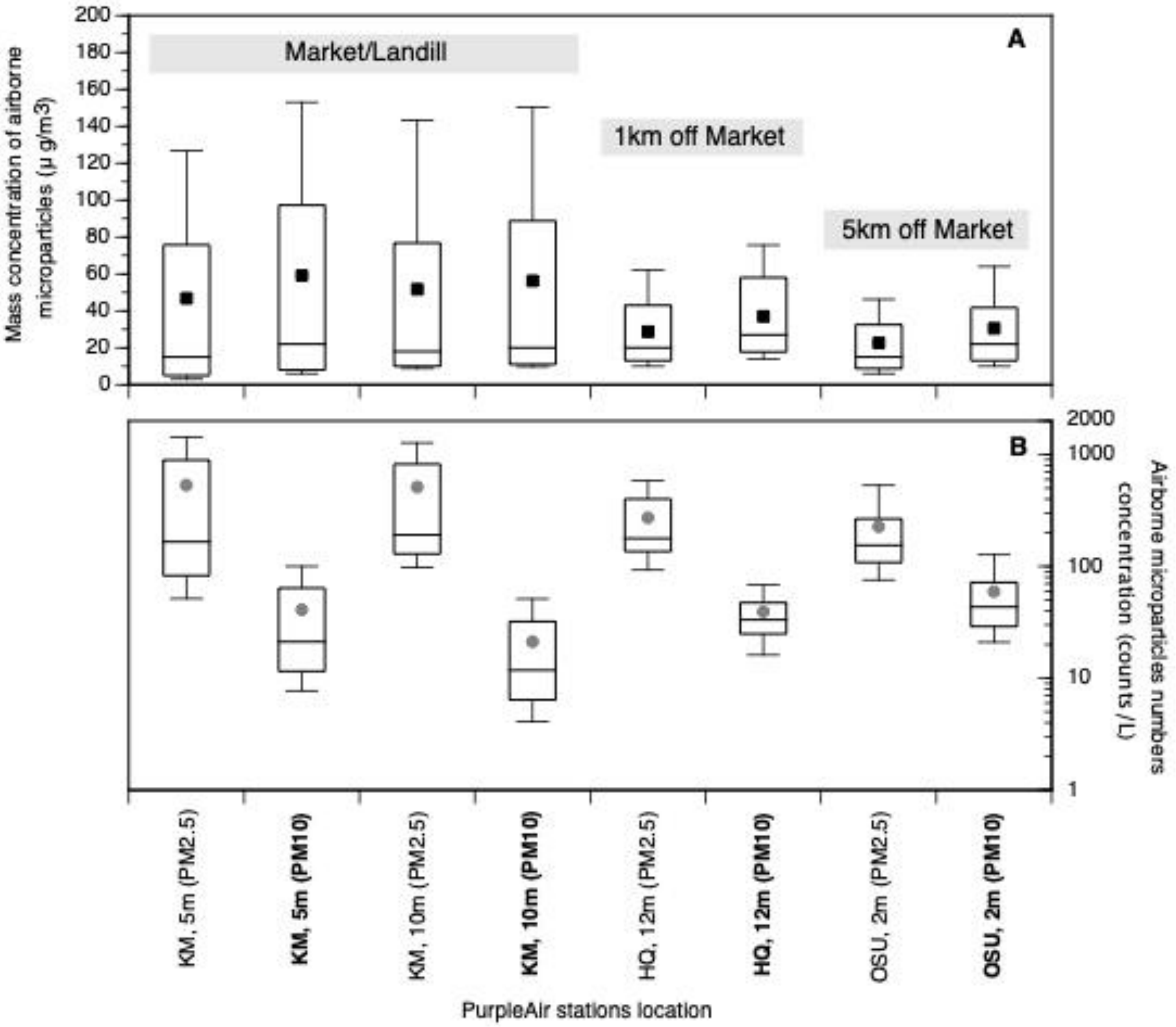

Online Resource 5

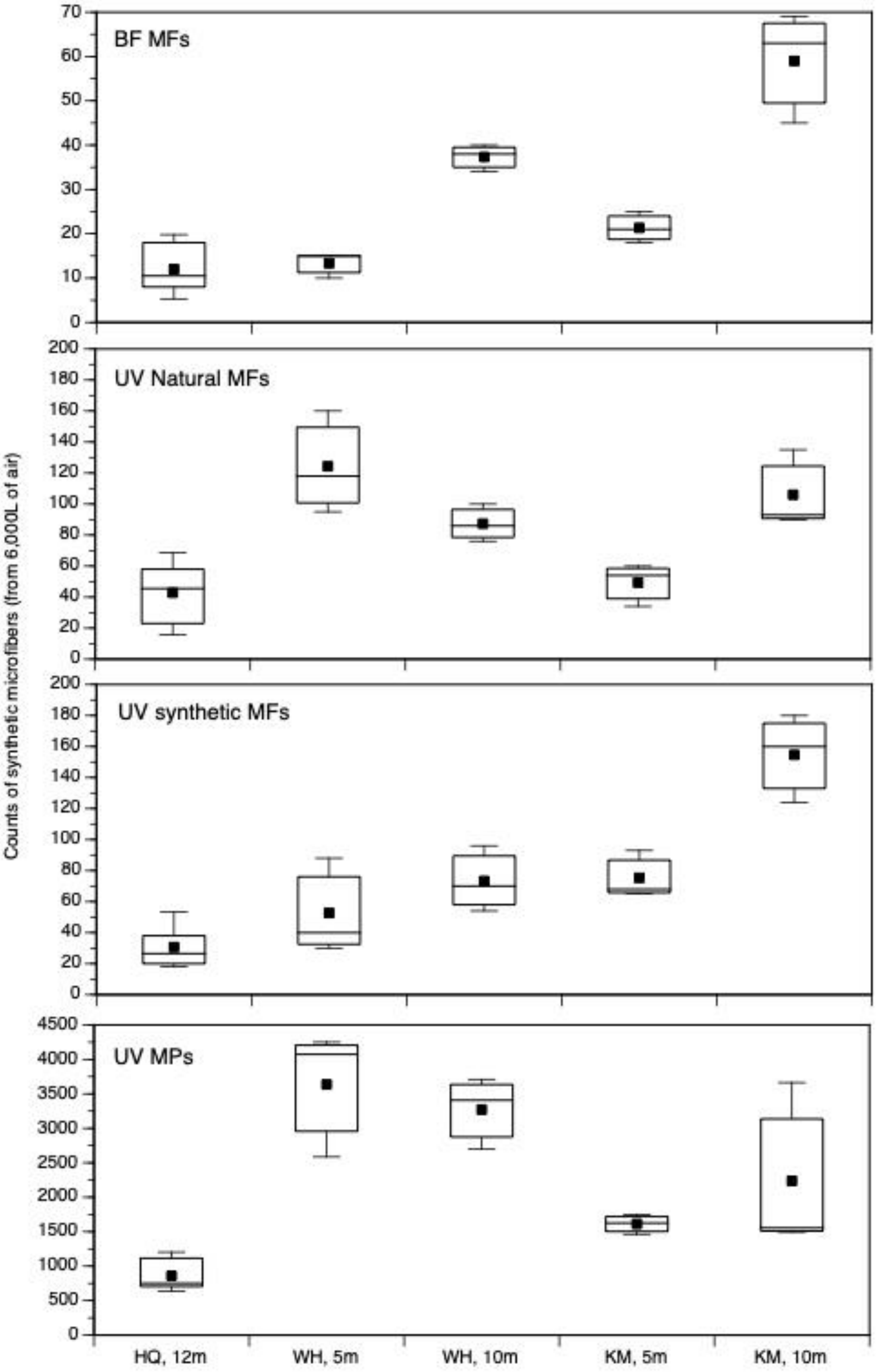


343

Online Resource 6

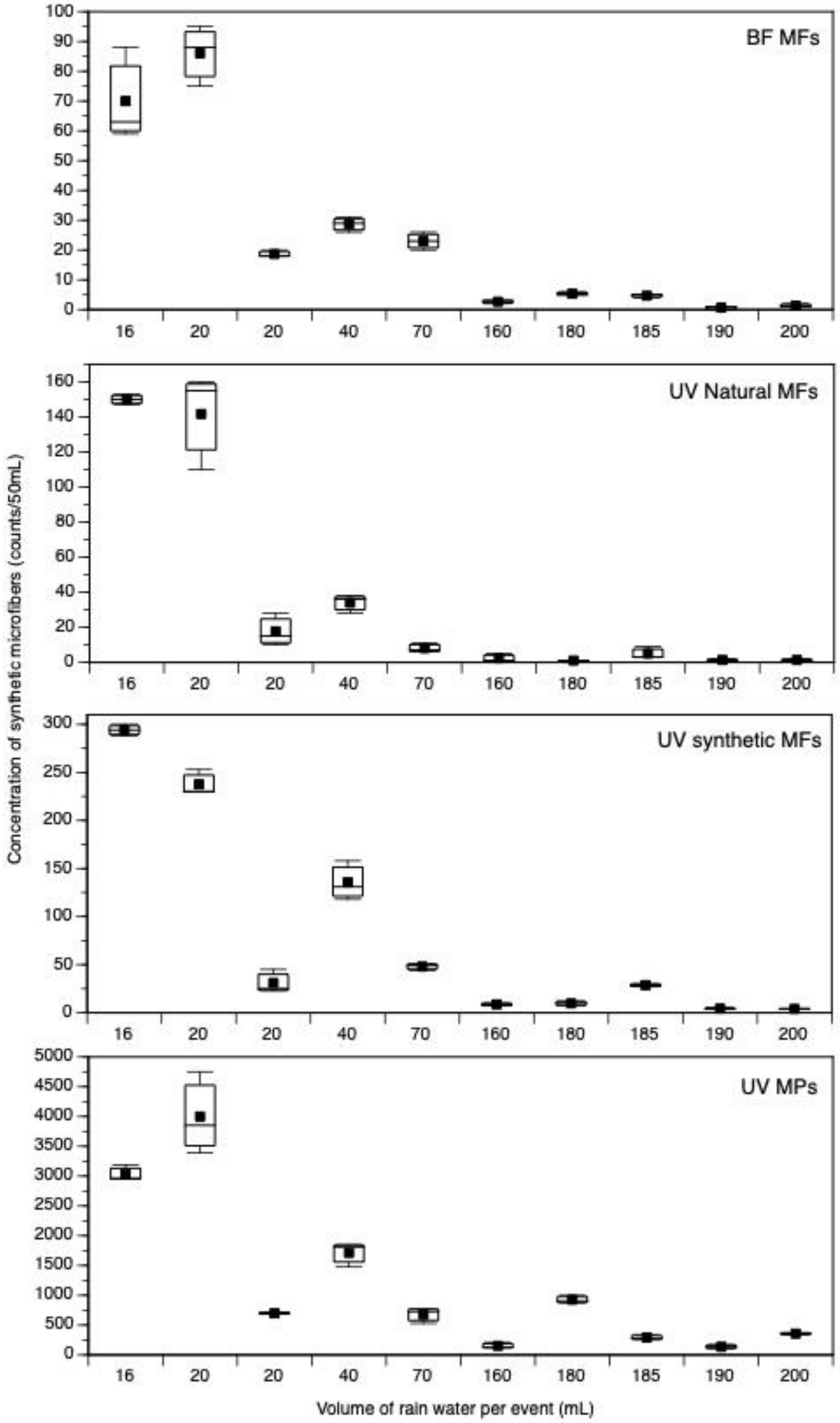

Online Resource 7

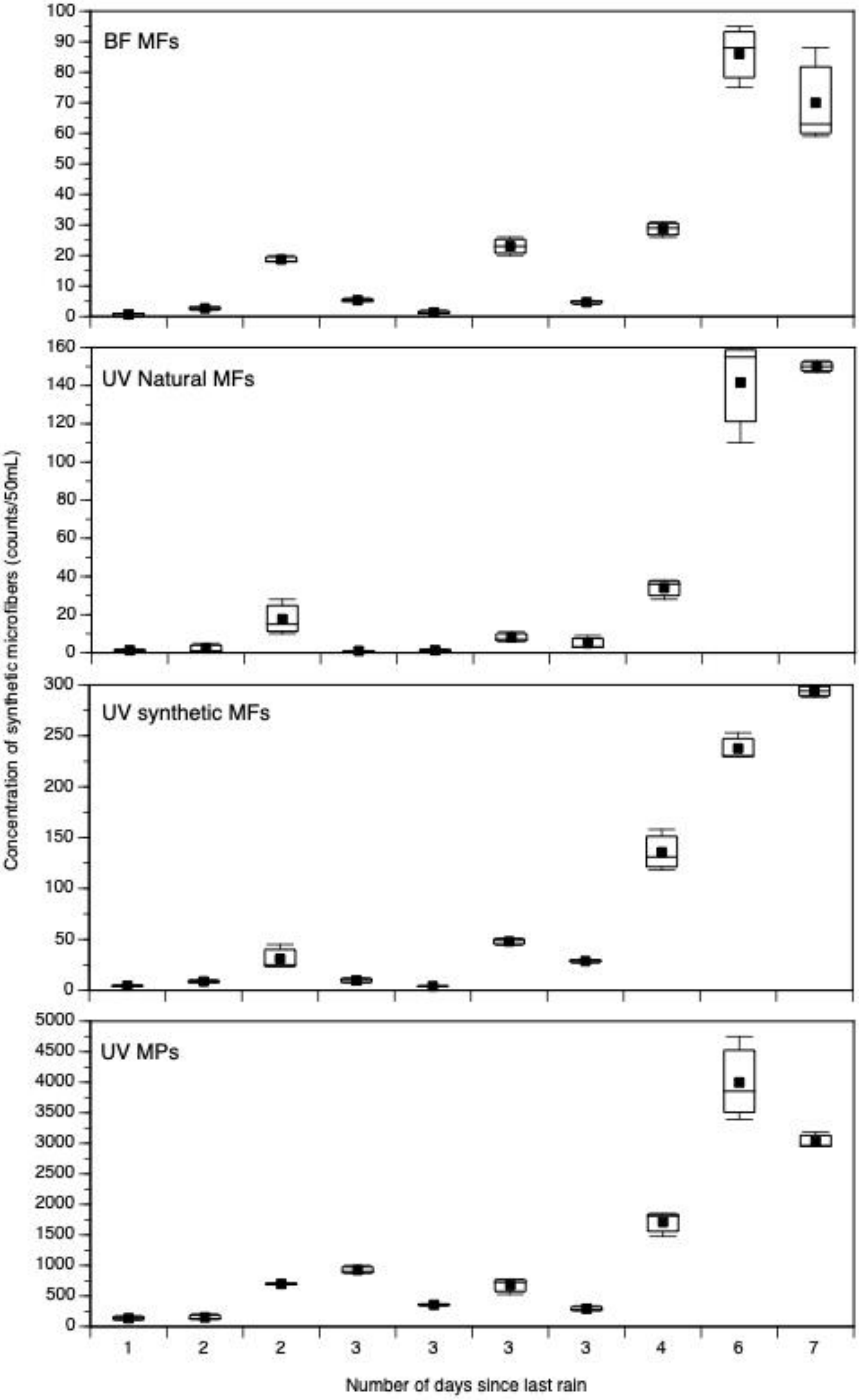


345

Online Resource 8

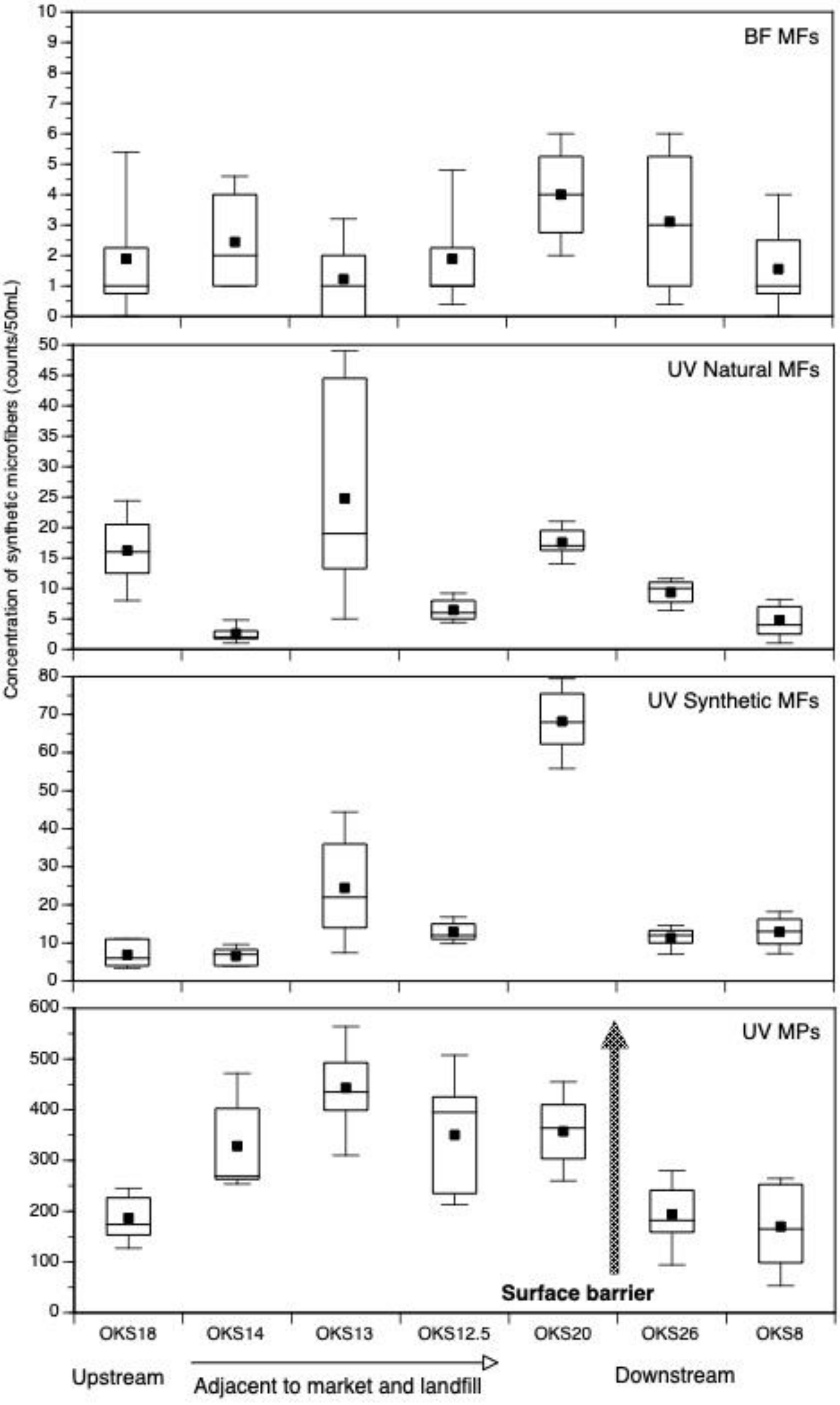


346

**Online Resource 9.** Minimum (Min), Maximum (Max), Mean and Median values for the counts of microfibers and microplastics identified as colored in bright field (freshly released) or with red fluorescence in epifluorescence (natural), from samples of air (6,000 L), rainwater (variable volumes), and Korle lagoon water (50 mL). For all counts, N=9, except for the airborne sampling counts (N=3). Abbreviations: HQ: HeadQuarter of The Or Foundation (roof) at two different times (a,b); WH: WareHouse space adjacent to the market at two different heights (5 m, 10 m); KM Kantamanto Market (center) at two different heights (5, 10 m). All lagoon sample counts originated from 50 mL of surface water, thus with concentration expressed in “counts/50 mL”, while airborne microparticle samples originated from 6,000 L of processed air, thus with “counts/6,000 L”.

| | | Bright field | | | | Red epifluorescence | | | |
|---|---|---|---|---|---|---|---|---|---|
| | | **Min** | **Max** | **Mean** | **Median** | **Min** | **Max** | **Mean** | **Median** |
| Airborne particles (counts/6,000 L) | **HQa** | 5 | 8 | 7 | 7 | 9 | 28 | 20 | 23 |
| | **HQb** | 2 | 3 | 3 | 3 | 6 | 19 | 14 | 17 |
| | **WH – 5 m** | 4 | 6 | 5 | 6 | 38 | 64 | 50 | 47 |
| | **WH – 10 m** | 17 | 20 | 19 | 19 | 38 | 50 | 44 | 43 |
| | **KM – 5 m** | 14 | 20 | 17 | 17 | 27 | 48 | 39 | 43 |
| | **KM – 10 m** | 36 | 55 | 47 | 50 | 72 | 108 | 85 | 74 |
| Rainwater (HQ) | **16 mL** | 19 | 28 | 22 | 20 | 47 | 49 | 48 | 48 |
| | **20 mL** | 30 | 38 | 34 | 35 | 44 | 64 | 57 | 62 |
| | **160 mL** | 7 | 9 | 8 | 9 | 2 | 16 | 7 | 3 |
| | **190 mL** | 1 | 2 | 2 | 2 | 4 | 7 | 5 | 5 |
| | **180 mL** | 17 | 20 | 18 | 17 | 2 | 4 | 3 | 2 |
| | **200 mL** | 3 | 6 | 5 | 5 | 3 | 6 | 5 | 5 |
| | **185 mL** | 16 | 19 | 18 | 18 | 10 | 33 | 18 | 12 |
| | **70 mL** | 28 | 36 | 32 | 32 | 9 | 16 | 12 | 10 |
| | **40 mL** | 21 | 25 | 23 | 23 | 22 | 30 | 27 | 29 |
| | **20 mL** | 7 | 8 | 7 | 7 | 4 | 11 | 7 | 6 |
| Korle lagoon sites (counts/50 mL) | **OKS18** | 0 | 7 | 2 | 1 | 6 | 26 | 16 | 16 |
| | **OKS14** | 1 | 5 | 2 | 2 | 1 | 6 | 3 | 2 |
| | **OKS13** | 0 | 4 | 1 | 1 | 5 | 51 | 25 | 19 |
| | **OKS12.5** | 0 | 6 | 2 | 1 | 4 | 10 | 6 | 6 |
| | **OKS20** | 2 | 6 | 4 | 4 | 14 | 21 | 18 | 17 |
| | **OKS26** | 0 | 6 | 3 | 3 | 6 | 12 | 9 | 10 |
| | **OKS8** | 0 | 4 | 2 | 1 | 1 | 9 | 5 | 4 |
